# YAP orchestrates heterotypic endothelial cell communication via HGF/c-MET signaling in liver tumorigenesis

**DOI:** 10.1101/2020.03.17.995225

**Authors:** Stefan Thomann, Sofia M. E. Weiler, Simone Marquard, Claudia R. Ball, Marcell Tóth, Teng Wei, Carsten Sticht, Carolina De La Torre, Eduard Ryschich, Olga Ermakova, Carolin Mogler, Daniel Kazdal, Norbert Gretz, Hanno Glimm, Eugen Rempel, Peter Schirmacher, Kai Breuhahn

## Abstract

Next to cell autonomous mechanisms, the oncogene *yes-associated protein* (YAP) controls liver tumor initiation and progression *via* cell extrinsic functions creating a tumor-supporting environment. However, how YAP affects the microenvironment and in particular the vascular niche, which contributes to liver disease and hepatocarcinogenesis, is poorly understood.

In this study, histo-morphological and molecular characterization of murine liver *endothelial cells* (ECs) populations and human single cell data revealed the presence of *liver sinusoidal endothelial cells* (LSECs) and *capillary endothelial cells* (CECs) in healthy liver tissue. In YAP^S127A^-induced tumorigenesis, a gradual replacement of LSECs by CECs was associated with dynamic changes in the expression of genes involved in EC subtype-specific paracrine communication. The formation of new potential communication hubs connecting CECs and LSECs included the *hepatocyte growth factor* (Hgf)/c-Met signaling pathway. In hepatocytes and tumor cells, YAP/*TEA domain transcription factor 4* (TEAD4)-dependent transcriptional induction of *osteopontin* (Opn) stimulated c-Met expression in ECs with CEC phenotype, which sensitized these cells to the pro-migratory effects of LSEC-derived Hgf. In human HCCs, the presence of a migration-associated tip-cell signature correlated with poor clinical outcome and the loss of LSEC marker gene expression. In addition, the replacement of LSECs by CECs with exclusive c-MET expression in a CEC subpopulation was confirmed at the single cell level.

In summary, YAP-dependent changes of the liver vascular niche comprise the formation of heterologous communication hubs (e.g. the HGF/c-Met axis), in which tumor cell-derived factors modify the crosstalk between LSECs and CECs.

## Introduction

The Hippo signaling pathway negatively controls the subcellular localization and activity of the transcriptional co-activator *yes-associated protein* (YAP). Indeed, inactivation of Hippo pathway constituents or nuclear enrichment of YAP has been associated with tumor formation as exemplified for many solid cancer entities such as *hepatocellular carcinoma* (HCC) (1).

So far, first tumor cell intrinsic mechanisms have been described for YAP that contribute to massive liver overgrowth and malignant transformation. These include dedifferentiation of liver hepatocytes associated with elevated proliferation and accumulation of oncogenic mutations (2,3). In addition, the Hippo/YAP axis transcriptionally controls the expression of genes critically involved in replication, chromosomal segregation, as well as DNA repair and dysregulation of these factors increases the risk of *chromosome instability* (CIN) (4–6). However, recent data suggested that next to tumor cell autonomous effector mechanisms, aberrant YAP activation in malignantly transformed cells influences immune cells to create a tumor-supportive environment. For example, YAP affects recruitment and polarization of myeloid-derived suppressor cells and immune evasion in an HCC cancer model (7). This data points to a connection between YAP activation in tumor cells and non-tumorous cells as well as in between non-malignant cells (e.g. *non-parenchymal* liver-resident *cells*; NPCs) *via* paracrine-acting growth factors, cytokines, and chemokines. Especially how YAP controls heterologous cell communication of vessel-forming *endothelial cells* (ECs) in liver tumorigenesis is not well understood.

The liver vascular network consists of phenotypically different ECs (8–10). Organotypic *liver sinusoidal endothelial cells* (LSECs) connect the vessels of the portal field with the central vein (porto-central axis) and are characterized by morphological features such as sieve plate fenestrations and the absence of a basement membrane. In contrast, portal and pericentral vessels are lined by a continuous endothelium (*capillary endothelial cells*; CECs), which is covered by an additional layer of smooth muscle cells in the portal field to withstand higher pressure and pulsatile flow. LSECs and CECs differ in their endocytotic capacity, antigen presentation (11), and marker gene expression (e.g. LSEC-specific Lyve-1, CD32b, Clec4g, Stabilin-2) (12). It was therefore hypothesized that EC subtypes may exert specialized functions with regard to liver development, metabolic zonation, and the regulation of the blood-flow rate (13,14). Clinically, dynamic alterations of LSECs and CECs are frequently observed in chronic liver disease and hepatocarcinogenesis. Here, the LSEC phenotype is progressively lost and replaced by non-fenestrated ECs - a process called capillarization (15,16). If and to which extent this process is controlled by YAP-mediated extrinsic functions derived from tumor cells is not known.

In this study, we aimed to dissect the dynamics in EC population abundance and its communication patterns in normal livers as well as YAP-induced hepatocarcinogenesis. Our data unravel central paracrine hubs connecting liver EC subtypes (e.g. *hepatocyte growth factor* (Hgf)/c-Met) as well as ECs and parenchymal cells (e.g. *osteopontin* (Opn)), which contribute to the formation of a tumor-supporting microenvironment.

## Materials and Methods

Antibodies used for western immunoblotting or immunohistochemical staining as well as corresponding dilutions, primer and siRNA sequences are listed in supplementary tables. A detailed description of the *in vivo* and *in vitro* analyses as well as protocols for mouse work, isolation of primary cells (CECs and LSECs) and sample preparation, real-time PCR, western immunoblotting, expression profiling, tissue-microarray analyses (including identifiers of data), and bioinformatics as well as statistical procedures can be found in supplementary materials and methods.

### Access to original expression profiling data for reviewers (3 data sets)

**Expression data of wildtype ECs (CECs and LSECs)**

https://www.ncbi.nlm.nih.gov/geo/query/acc.cgi?acc=GSE128042 Enter token azktgcmytbqnvqh into the box

**Dynamic expression data of CECs and LSECs (wt, early, and late)**

https://www.ncbi.nlm.nih.gov/geo/query/acc.cgi?acc=GSE128044 Enter token ipijmkciztehrkb into the box

**Expression data of SVEC4-10 after HGF stimulation (3 and 6 hrs)**

https://www.ncbi.nlm.nih.gov/geo/query/acc.cgi?acc=GSE128046 Enter token szwpwqsgfvsldwp into the box

## Results

### Morphological, functional, and molecular characterization of liver EC subpopulations

For a first characterization of EC diversity in mouse livers, we analyzed different surface marker combinations that enabled the discrimination of CECs and LSECs. By using the pan-EC markers CD146 and CD31 in combination with the sinusoidal-specific markers Lyve-1 and CD32b, both cell-types were clearly separated using tissue *immunofluorescence* (IF). While CECs lining the vessels in the portal tract and the central vein were CD146^+^/Lyve-1^−^(CD146^+^/CD32b^−^, CD31^+^/Lyve-1^−^, CD31^+^/CD32b^−^), sinusoidal LSECs were CD146^+^/Lyve-1^+^ (CD146^+^/CD32b^+^, CD31^+^/Lyve-1^+^, CD31^+^/CD32b^+^), (Figure 1A, Suppl. Figure S1A). The validity of this histo-morphological approach was illustrated by fluorescence intensity-based plotting of porto-sinusoidal transition zones, showing a continuous CD146 positivity in all EC subtypes, while Lyve-1 expression was restricted to LSECs (Figure 1B). Quantitative measurements of multiple porto-sinusoidal transition zones illustrated a consistent expression of CD146 in porto-sinusoidal transition zones, while Lyve-1 was exclusively detectable in sinusoids (Suppl. Figure S1B).

**Figure 1:**
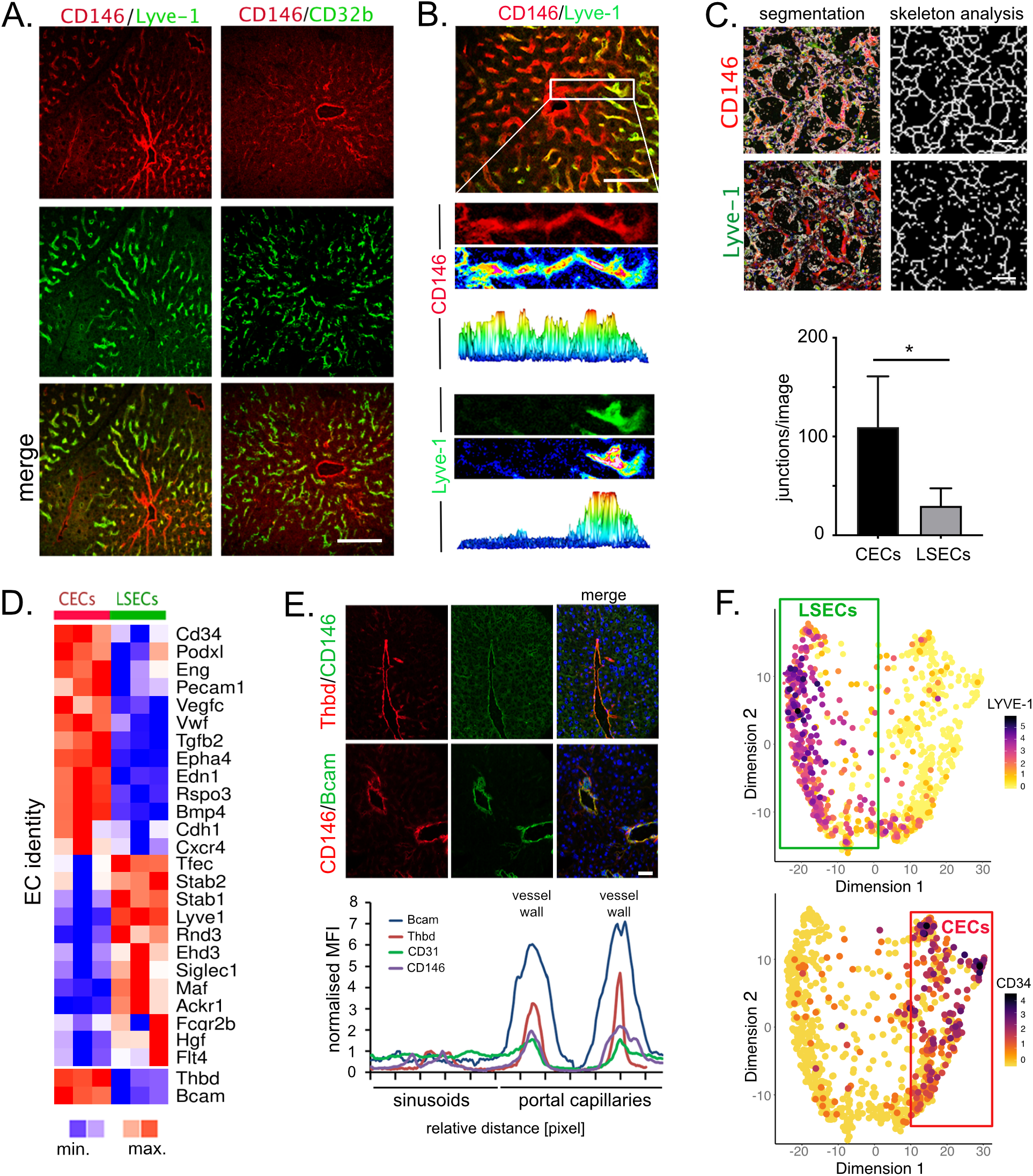
Dissection of liver EC subtypes. **(A.)** CD146 (red) and CD32b/Lyve-1 (green) double IF stains to characterize liver-resident EC populations: CECs (CD146^++^/Lyve-1^−^or CD146^++^/CD32b^−^) and LSECs (CD146^+^/Lyve-1^+^ or CD146^+^/CD32b^+^). Scale bar: 50 μm. **(B.)** CD146 (red) and Lyve-1 (green) double IF stains followed by intensity measurements of porto-sinusoidal transition zones. Scale bar: 50 μm. **(C.)** Modified tube formation assay of primary heterogenous liver ECs followed by IF with CD146 (red) and Lyve-1 (green). After segmentation of CEC-(CD146^+^/Lyve1^−^) and LSEC-(CD146^+^/Lyve-1^+^) specific networks, junctions were automatically quantified. ECs were cultured up to 5 days to allow endothelial network formation. Graphs show mean ± SD (12 images per group); Mann-Whitney U test (*p≤0.05). Scale bar: 50 μm. **(D.)** Heatmap of 27 differentially regulated genes selected and used as LSEC and CEC denominators. Alongside known genes (e.g. Stab2 for LSECs) several new potential subtype marker genes are identified (e.g. Bcam, Thbd). FDR≤0.05 (n=3). **(E.)** Bcam (green)/CD146 (red) and Thbd (red)/CD146 (green) double IF stains of larger liver vessels. Scale bar: 50 μm. Normalized and marker-specific MFI profiles of peri-portal and portal liver tissues. Exemplary MFI values of pan-EC markers CD31 and CD146 as well as CEC-selective markers Bcam and Thbd are shown. Two peaks in the portal tract indicate CEC vessel walls. **(F.)** ScRNA-seq data derived from nine healthy human liver samples were re-analyzed (19). Only cells from clusters 9, 10, 13, 20, 29, and 32 expressing PECAM1/CD31 were chosen for *PHATE* visualization. Exemplary, LYVE-1 as LSEC marker and CD34 as CEC marker are shown.

Isolated ECs exhibited the typical cobblestone morphology in bright field microscopy and actively took up *acetylated low-density lipoprotein* (AcLDL), (Suppl. Figure S1C). IF and FACS analysis for CD146 and Lyve-1 illustrated a mixed EC cell-population including CD146^+^/Lyve-1^−^(CECs; 4±2%) and CD146^+^/Lyve-1^+^ cells (LSECs; 95±2%), (Suppl. Figure S1D). Scanning electron microscopy confirmed the presence of LSEC-specific sieve plate fenestrations in most isolated cells, while ECs without fenestrations represented CECs (Suppl. Figure S1E). To study functional differences between CECs and LSECs, a capillarization-inducing cultivation protocol was used to analyze network formation of both EC subtypes (17). Indeed, IF stains for CD146/Lyve-1 followed by comparative analysis of EC networks illustrated that CECs formed significantly more junction points than LSECs (Figure 1C).

One requirement for a molecular characterization of CECs and LSECs was the isolation of pure cell fractions. For this, MACS-based pre-purified CD146^+^ ECs were sorted for CD146, CD31, and Lyve-1 using a FACS-based protocol (Suppl. Figure S2A/B). The approach was confirmed by real-time PCR using CEC markers (Vwf, Edn1) and LSEC markers (Lyve-1 and *c-type lectin domain family member b* (Clec1b)), (Suppl. Figure S2C) (18).

Comprehensive transcriptome analysis revealed that 247 genes were differentially expressed between CECs and LSECs (Suppl. Figure S2D; FDR≤0.05). As expected, genes that characterized EC identity including *Stabilin1/2*, *CD34*, *MAF BZIP transcription factor* (Maf), and *transcription factor EC* (Tfec) were differentially expressed between both EC subtypes (Figure 1D). *Gene set enrichment analysis* (GSEA) also revealed an accumulation of differentially expressed genes involved in Hippo- and *transforming growth factor-beta* (TGFβ)-signaling in CECs (Suppl. Figure S2E). In addition, CECs showed positive enrichments of gene sets involved in cell mobility (e.g. Itgb4, Itga2), cell adhesion (e.g. Ncam1, L1cam), and cellular guidance (e.g. Sema3f, Sema3d), suggesting that CECs exhibit the molecular requirements for active migration.

Several genes differentially expressed between CECs and LSECs have not been described as potential subtype markers before. We selected *basal cell adhesion molecule* (*Bcam*) and *thrombomodulin* (*Thbd/CD141*), which were highly expressed in CECs but not in LSECs (Figure 1D/E). Real-time PCR analysis as well as IF stains illustrated highest signal intensity for Bcam and Thbd in CECs of the of the portal tracts but not in LSECs in sinusoids (Figure 2E, Suppl. Figure S2F). MFI measurements revealed that the relative intensities of Bcam and Thbd were higher in vessels of the portal tract than CD146 or CD31, suggesting that these proteins may represent highly specific markers for the discrimination of discontinuous and continuous liver ECs (Figure 1E, graph).

**Figure 2:**
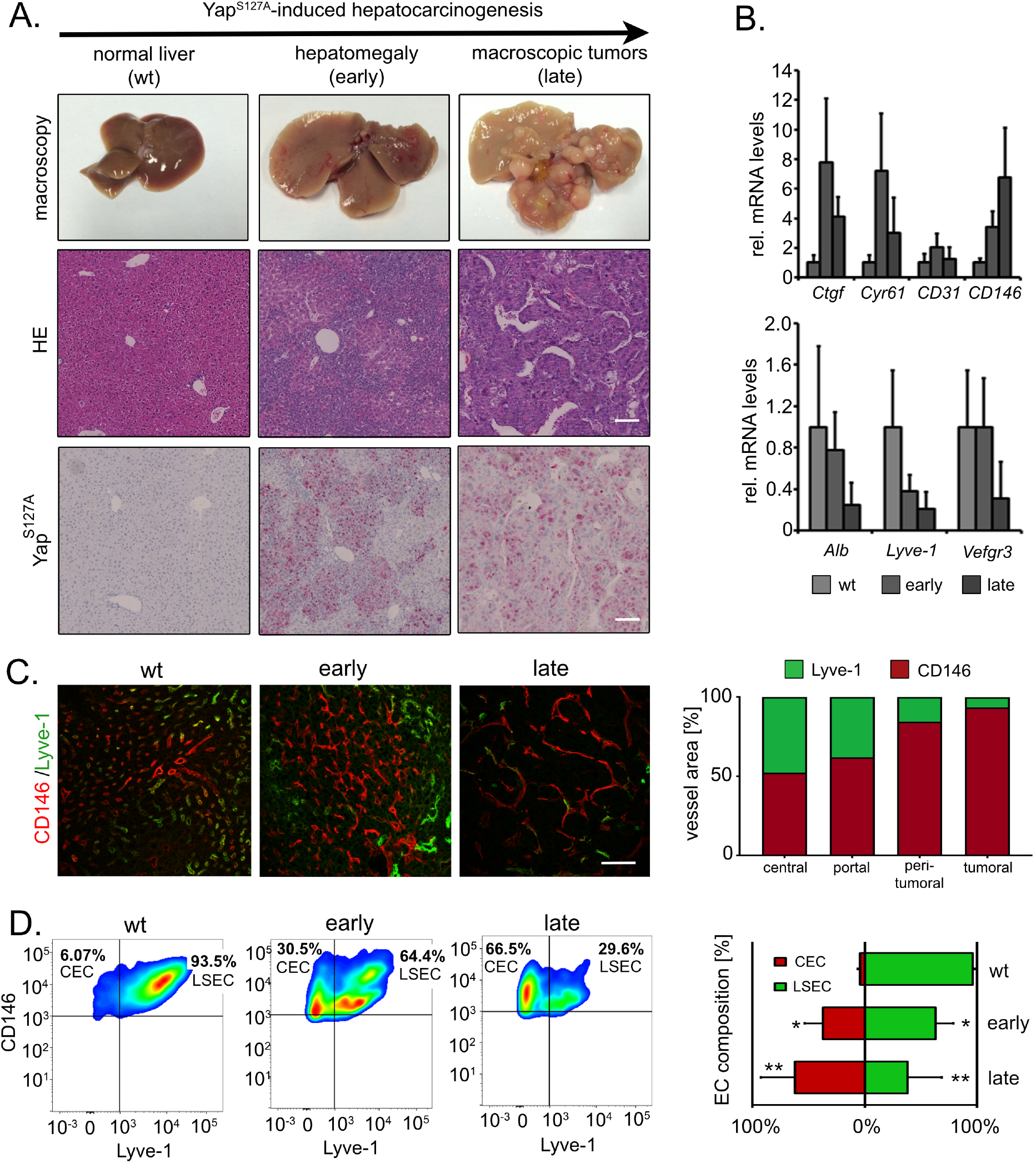
EC class switch in premalignant and malignant mouse livers. **(A.)** Induction of constitutively active Yap^S127A^ is leads to hepatomegaly and tumor formation. Histological H&E sections illustrating the expansion of densely-packed hepatocellular cells in hepatomegaly (early) and tumor-bearing livers (late). Immunohistochemistry reveals Yap positivity in mitotically active tissue areas. Scale bar: 100 μm. **(B.)** Real-time PCR analysis of Yap target genes (Ctgf, Cyr61), a marker of hepatocellular differentiation (albumin, Alb), and EC markers (CD31, CD146, Lyve-1, and Vegfr3). Light grey bars: normal livers (wt); grey bars: hyperplastic (early) livers; dark grey bars: tumor-bearing (late) livers. Graphs show mean ± SD (wt n=10; early n=4; late n=11). **(C.)** CD146 (red) and Lyve-1 (green) IF stains were subjected to confocal microscopy and image segmentation to discriminate between CEC-(red) and LSEC-specific (green) vessel areas. Red bar: percentage of CD146-positive vessel area; green bar: percentage of Lyve-1-positive vessel area (relative to total image area). Graphs show CD146 or Lyve-1 vessel areas. In total, 12-13 images/group were analyzed (4-5 images/mouse, n=3). Scale bar: 50 μm. **(D.)** Exemplary FACS analysis of CEC and LSEC composition in normal livers as well as early and late stages of oncogene-induced tumorigenesis. Red bars: percentage of CEC population; green bars: percentage of LSEC population. Graphs show mean ± SD (n=4-8/group). ANOVA and post hoc testing, wt *vs.* early *p≤0.05; wt *vs.* late **p≤0.01.

Lastly, validity of the sorting approach followed by bulk analysis was confirmed using single-cell RNA sequencing (scRNA-seq) data derived from healthy human liver tissues (19). These results illustrated the presence of two distinct subgroups of ECs, which are characterized by the presence CEC markers (e.g. CD34, EPHA4, VWF, and BCAM) or LSEC markers (e.g. LYVE-1, STAB2, FLT4, and CLEC4G), (Figure 1F, Suppl. Figure S3).

In summary, these results demonstrate the existence of two EC subtypes in mouse and human liver with distinct histo-morphological, functional, and molecular characteristics.

### Dynamic changes of EC composition in YAP-induced hepatocarcinogenesis

In the next step we were interested if dysregulation of the oncogene YAP in hepatocytes could affect the vessel network at different stages of tumorigenesis. To approach these potential changes experimentally, we used the LAP-tTA/Yap^S127A^ transgenic mouse model for the inducible and hepatocyte-specific expression of constitutively active *yes-associated protein* (Yap^S127A^), which allows the investigation of EC subtypes under normal, premalignant, and malignant conditions without interfering fibrosis and inflammation (2).

Yap^S127A^-induced hepatomegaly and tumor formation within 8-15 weeks of transgene induction was associated with the expansion of Yap-positive and mitotically active cells (Figure 2A) (2,5). With regard to the expression of EC markers, tissue-based real-time PCR revealed an induction of Yap target genes (Ctgf, Cyr61) together with expression changes of EC markers (e.g. reduced LSEC markers Lyve-1/Vegfr3) in the process of YAP^S127A^-induced liver cancer development, implicating alterations of the EC composition (Figure 2B). Indeed, CD146/Lyve-1 IF stains revealed that Lyve-1 positivity decreased in peri- and intra-tumoral areas of Yap^S127A^-transgenic livers in comparison to porto-sinusoidal and sinusoidal-central transition zones of wildtype (wt) mice (Figure 2C).

Finally, dynamic changes within the EC population were quantitatively examined in normal, hyperplastic, and tumor-bearing livers by FACS analysis. The results demonstrated a significant reduction of LSECs from 95±2% in wt livers to 39±30% in late tumor stages, while the number of CEC increased from 4±2% in wt livers to 61±30% in tumors (Figure 2D). The model-independent shift from a LSEC to a CEC-phenotype in tumor tissue was confirmed in YAP-overexpressing autochthonous (AlbTag) and implantation (Hep56.1D) liver cancer models, suggesting that the EC population switch represents a model-independent process in hepatocarcinogenesis (Suppl. Figure S4A/S4B).

These results unveil a progressive replacement of LSECs by CECs in YAP-induced hepatocarcinogenesis, which recapitulates a phenotype observed in human chronic liver disease and cancer.

### CECs and LSECs evolve distinct expression signatures in YAP-dependent tumorigenesis

Our previous results demonstrated an EC subpopulation shift in YAP-driven hepatocarcinogenesis; however, it is unknown to what extent CECs and LSECs dynamically modify their gene expression in this process *in vivo*. To define their dynamic molecular response in tumorigenesis, CECs and LSECs were isolated from wt as well as Yap^S127A^-induced hyperplastic (early) and tumor-bearing (late) livers. A comparative analysis of CEC- and LSEC-specific marker gene expression validated the purity of both EC populations at all stages of the experiment (Figure 3A).

**Figure 3:**
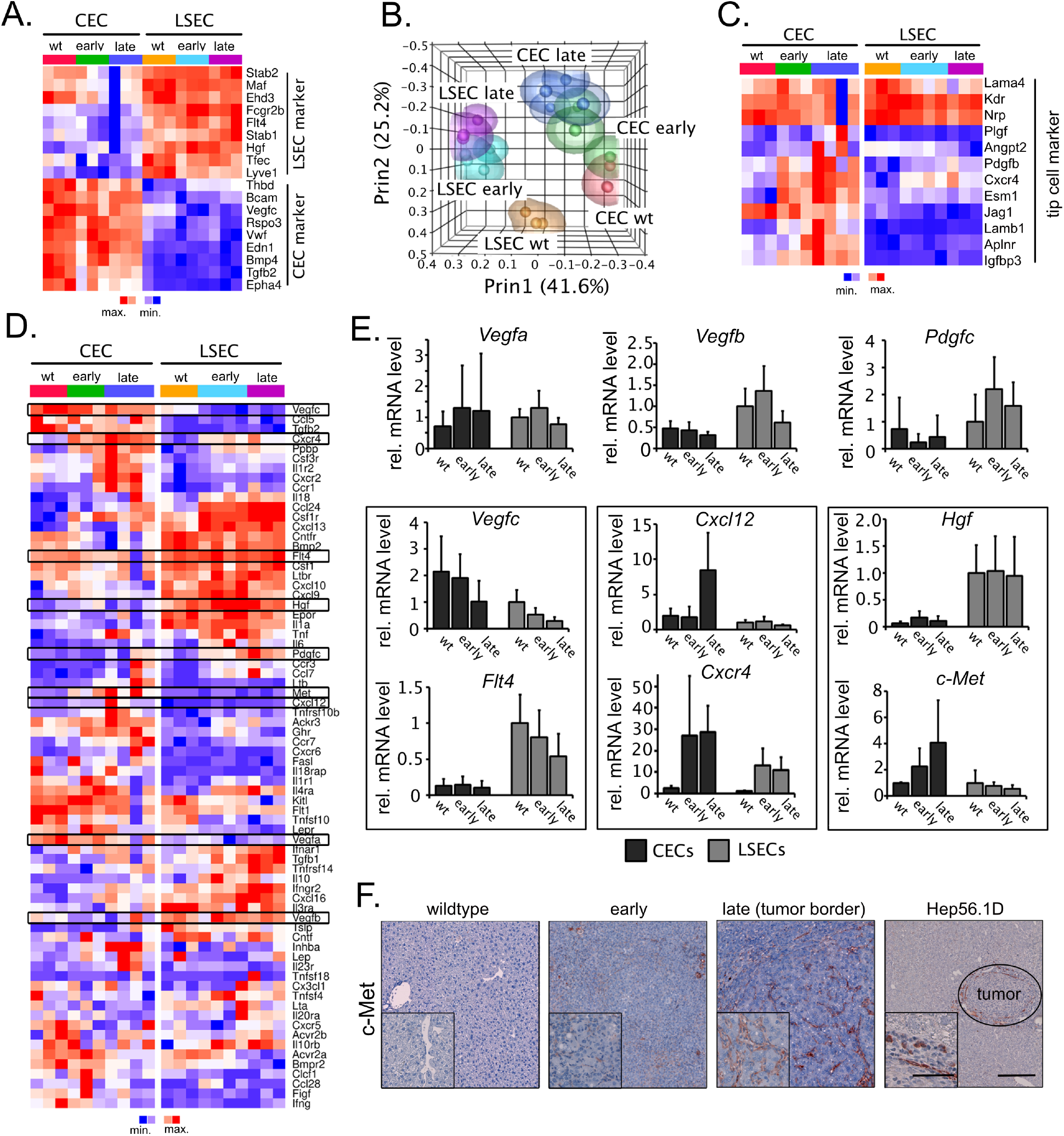
CEC- and LSEC-specific gene expression patterns in liver tumorigenesis. **(A.)** Expression of 9 CEC and 9 LSEC markers in sorted CEC and LSEC subgroups isolated from mice with wt, hyperplastic (early) and tumor-bearing (late) livers illustrate sorting efficiency of samples used for transcriptome analysis, (n=3). For all further analyses, an additional sample for CEC (late) and LSEC (early) was included (n=4). **(B.)** PCA visualizes transcriptome differences between CECs and LSECs as well as their evolution related to the disease stage (n=3-4). **(C.)** Heatmap illustrating the dynamic expression of 12 tip cell markers in CECs and LSECs (n=3-4). **(D.)** Heatmap visualizing the differential and dynamic expression of significantly regulated genes in CECs and LSECs listed in ‘cytokine-cytokine receptor interaction’ (KEGG: mmu04060). Additional paracrine-acting genes were included (Vegf family: Vegfa, Vegfb, Vegfc, Figf (Vegfd), Flt1 (Vegfr1), Flt3 (Vegfr3) and Pdgfc, Hgf, c-Met). Genes differentially expressed between EC subgroups (e.g. CEC wt *vs.* LSEC wt) or between disease stages (e.g. LSEC wt *vs.* LSEC early) were considered for this analysis (n=3-4). FDR≤0.05. **(E.)** Real-time PCR confirmation of 9 genes highlighted in (D.). Black bars: CECs; grey bars: LSECs. Known ligand/receptor combinations are highlighted by black boxes (n=3-4). **(F.)** Immunohistochemical stains of c-Met in wt, hyperplastic and tumor-bearing livers derived from Yap^S127A^-transgenic animals and from an orthotopic implantation model (Hep56.1D). No obvious c-Met positivity of hepatocytes or tumor cells is detectable. Scale bar (lower magnification): 200 μm; scale bar (higher magnification): 50 μm.

In total, 4,877 distinct genes with KEGG pathway annotation were significantly regulated when multiple comparisons between CECs and LSECs as well as between EC subtypes at different stages of tumorigenesis were performed (Suppl. Figure S5A). *Principal component analysis* (PCA) illustrated highly concordant gene expression within all six biological groups (CEC- and LSEC-specific clusters) as well as the molecular evolution of both cell-types within carcinogenesis (Figure 3B). To identify CECs- and LSEC-specific properties in tumor angiogenesis, a panel of 11 published genes that characterize highly polarized cells within angiogenic sprouts was tested (tip cell signature) (20–22). Most genes of the signature were highly expressed in CECs or increased from wt to late stage CECs, which was indicative of a higher degree of cell migration in this EC subtype (Figure 3C). In accordance with these results, a differential expression of genes in the migration-associated KEGG pathways ‘focal adhesion’, and ‘cell adhesion and ECM interaction’ was detected (Suppl. Figure S5B).

Since we hypothesized that YAP dysregulation in hepatocytes affects the vascular compartment in a growth factor/cytokine/chemokine-dependent manner, YAP^S127A^-transgenic mice were treated with the FDA-approved multi-kinase inhibitor cabozantinib to block paracrine communication. Indeed, cabozantinib treatment abolished CEC expansion and reduced the expression of tip cell signature genes in YAP^S127A^-transgenic mice, confirming the presence of functionally relevant communication networks between different cells types such as EC subpopulations (Suppl. Figure S5C-S5E).

To identify potential dynamic cross-talks connecting CECs and LSECs subtypes in YAP^S127A^-induced tumor formation, only the dynamic behavior of genes involved in cellular signaling including ligands and receptors were compared (Figure 3D). Next to static, cell-type-specific differences (e.g. *transforming growth factor beta 2* (Tgfβ2) in CECs and *C-X-C motif chemokine ligand 9* (Cxcl9) in LSECs), real-time PCR analyses confirmed that several known ligand/receptor pairs were dynamically regulated in both EC subtypes. This was indicative of the existence of heterologous communication networks connecting CECs and LSECs in tumorigenesis (Figure 3E). These cytokine/cytokine receptor pairs included *C-X-C motif chemokine ligand 12* (Cxcl12)/*C-X-C motif chemokine receptor 4* (Cxcr4) and *vascular endothelial growth factor C* (Vegfc)/*fms related tyrosine kinase 4* (Flt4/Vegfr3). Equally, Hgf was highly expressed only by LSECs in wt, early, and late phases of liver tumorigenesis, while its receptor c-Met was progressively and exclusively induced in CECs (Figure 3E). This step-wise and CEC-specific induction of c-Met in tumorigenesis was confirmed in tissue specimens from Yap^S127A^ transgenic mice and in the orthotopic Hep56.1D implantation model (Figure 3F).

This data demonstrates that CECs and LSECs actively adjust their expression at the molecular level to form potential communication hubs in the process of YAP-dependent tumorigenesis.

### In YAP-induced carcinogenesis the Hgf/c-Met pathway controls CEC migration and invasiveness

To confirm the cell-type-specific expression of Hgf and c-Met in EC subtypes in an alternative model, a time-dependent differentiation culture model for primary mouse ECs was established (18). Here, short-term cultivation of ECs was characterized by a LSEC phenotype (≤ 2 days), while long-term cultured ECs lost the LSEC- and acquired the CEC phenotype (5 days). Indeed, a shift from an LSEC to a CEC phenotype after EC cultivation for 5-day was confirmed by an increased expression of different CEC markers (Suppl. Figure S6A). More importantly, Hgf was clearly reduced, while c-Met increased in cells with an CEC phenotype (Figure 4A). Lastly, Hgf levels in cell culture supernatants illustrated that freshly isolated ECs with an LSEC phenotype secreted drastically higher Hgf amounts than long-term cultured ECs with a CEC phenotype (Figure 4B).

**Figure 4:**
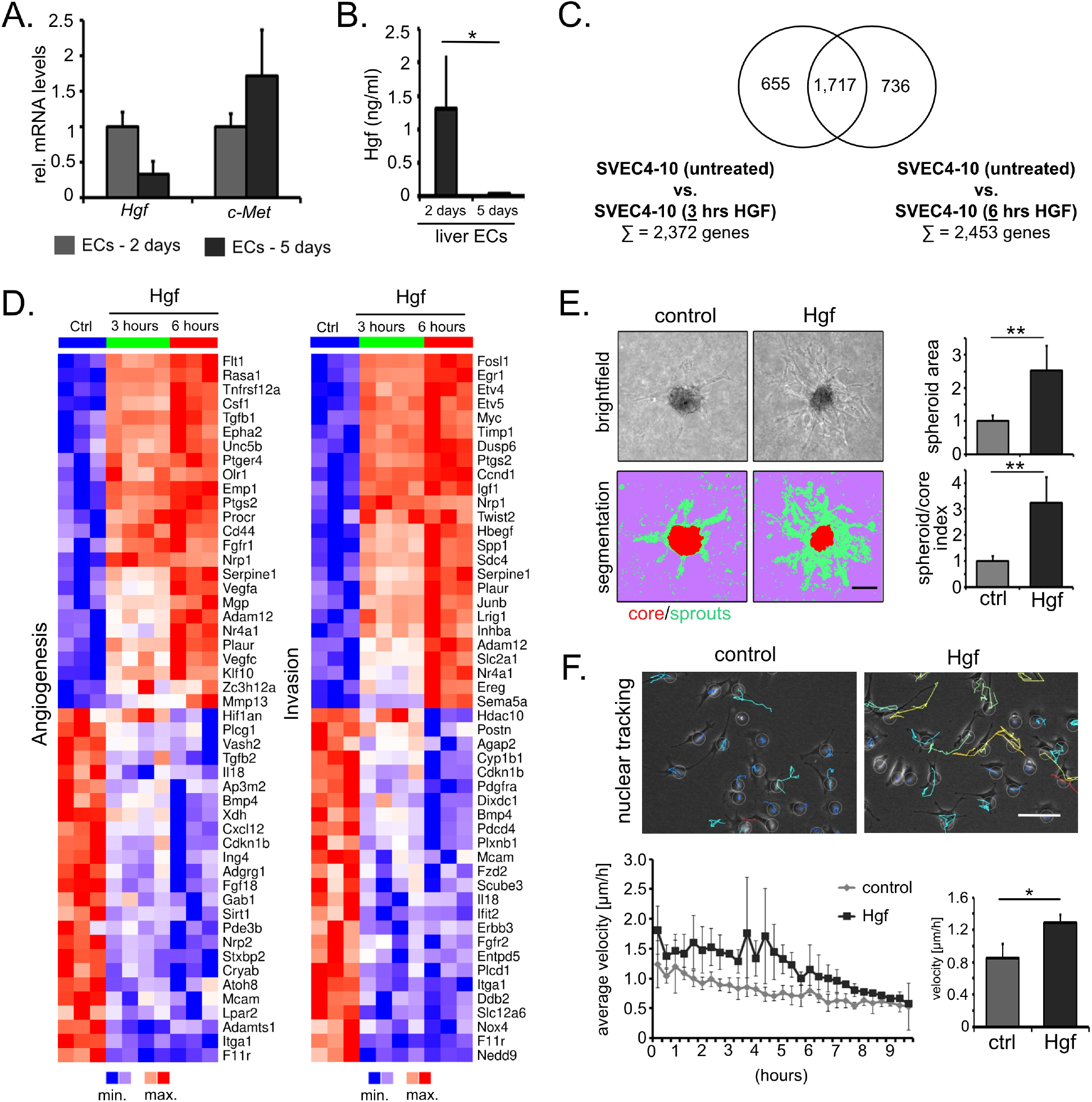
HGF stimulates cell migration in ECs with CEC characteristics. **(A.)** Real-time PCR analysis of *Hgf* and *c-Met* transcript levels in primary isolated murine ECs cultured for 2 days (grey bars, predominant LSEC phenotype) and 5 days (black bars, predominant CEC phenotype). Graphs show mean ± SD (n=4). Mann-Whitney U test (*p≤0.05). **(B.)** ELISA of secreted Hgf in the supernatant of ECs that were cultured for 2 or 5 days. Graph show mean ± SD (n=3-4/group). Mann-Whitney U test (**p≤0.01). **(C.)** Venn diagram summarizing significantly regulated genes in SVEC4-10 cells 3 and 6 hours after HGF stimulation (compared to untreated controls); n=3-4/group. Genes regulated in both groups were subjected to functional annotation clustering. **(D.)** Heatmaps showing the 50 strongest regulated genes (25 up and 25 down-regulated) in SVEC4-10 cells after Hgf stimulation for the process ‘angiogenesis’ and for the GO-ID (biological process) pathway ‘invasion’ (n=3-4); FDR p≤0.05. **(E.)** Cell-population-based spheroid assay of SVEC4-10 cells with and without Hgf stimulation. Sprouting was digitally documented after 48 hours. Graphs show mean ± SD (5-6 images/group). Mann-Whitney U test (**p≤0.01); one representative experiment is shown (total n=3). Scale bar: 50 μm. **(F.)** Time-lapse microscopy-based single cell tracking of SVEC4-10 cells with and without Hgf. Motion trajectories were documented for up to 10 hours. Colored lines in photographs illustrate motion trajectories. Time-dependent velocity of one representative experiment is shown. Graph on the right side shows mean ± SD (73.6 ± 22.9 cells/per visual field, 4 visual fields per experiment, n=4; *p≤0.05). Scale bar: 50 μm.

To further define the biological relevance of Hgf on CECs *in vitro* and because primary CECs are not suitable for functional experiments, the SV40-transformed cell line SVEC4-10 with CEC characteristics was used (23). SVEC4-10 cells express c-Met, lack the expression of typical LSEC markers (e.g. Lyve-1), and served as an Hgf-sensitive CEC-related model system (Suppl. Figure S6B/S6C). Expression profiling revealed that HGF administration for 3 and 6 hours led to the significant regulation of 1,717 genes (Figure 4C). *Gene ontology* (GO) analysis showed a regulation of genes involved in processes associated with blood vessel development (e.g. ‘angiogenesis’) and cell mobility (e.g. ‘invasion’), (Figure 4D, Suppl. Figure S6D). Together with our previous finding that CECs and LSECs differentially express cell mobility-associated genes (Suppl. Figures S2E and S5B), we hypothesized that the paracrine crosstalk between LSECs and CECs *via* Hgf/c-Met predominantly affects CEC migration.

Indeed, SVEC4-10 cell-population-based, 3-dimensional spheroid sprouting assays illustrated that Hgf significantly increased total spheroid size through the enhanced formation of sprouts in a surrounding matrix (Figure 4E). To substantiate the impact of Hgf on a single cell level, motion trajectories of SVEC4-10 cells were quantified after HGF administration using time-lapse microscopy. These results confirmed the previous cell-population-based data illustrating that Hgf stimulated higher cell velocity in comparison to untreated cells (Figure 4F).

In sum, this data demonstrates that in YAP-dependent tumor formation high-level secretion of Hgf by LSECs activates migration/invasion in ECs with a CEC phenotype.

### Endothelial responses are initiated by YAP-positive hepatocytes

Our data pointed to a dynamic communication between EC subtypes in the process of YAP-dependent pre-malignant liver capillarization and hepatocarcinogenesis. In order to describe potential mechanisms how hepatocytes and tumor cells initiate these changes in the vascular niche, we aimed to identify paracrine acting factors actively secreted by YAP^S127A^-positive parenchymal cells. Proteome array analysis revealed that several cytokines in conditioned medium from Yap^S127A^ transgenic hepatocytes were elevated, such as *osteopontin* (Opn), *placental growth factor* (Plgf), and *serpin family E member 1* (SerpinE1; syn. *plasminogen activator inhibitor*, Pai1), (Figure 5A/B, Suppl. Figure S7A). Indeed, confirmatory experiments illustrated high-level expression of Opn in primary hepatocytes at the mRNA level and in plasma of Yap^S127A^-transgenic mice (Figure 5C/D).

**Figure 5:**
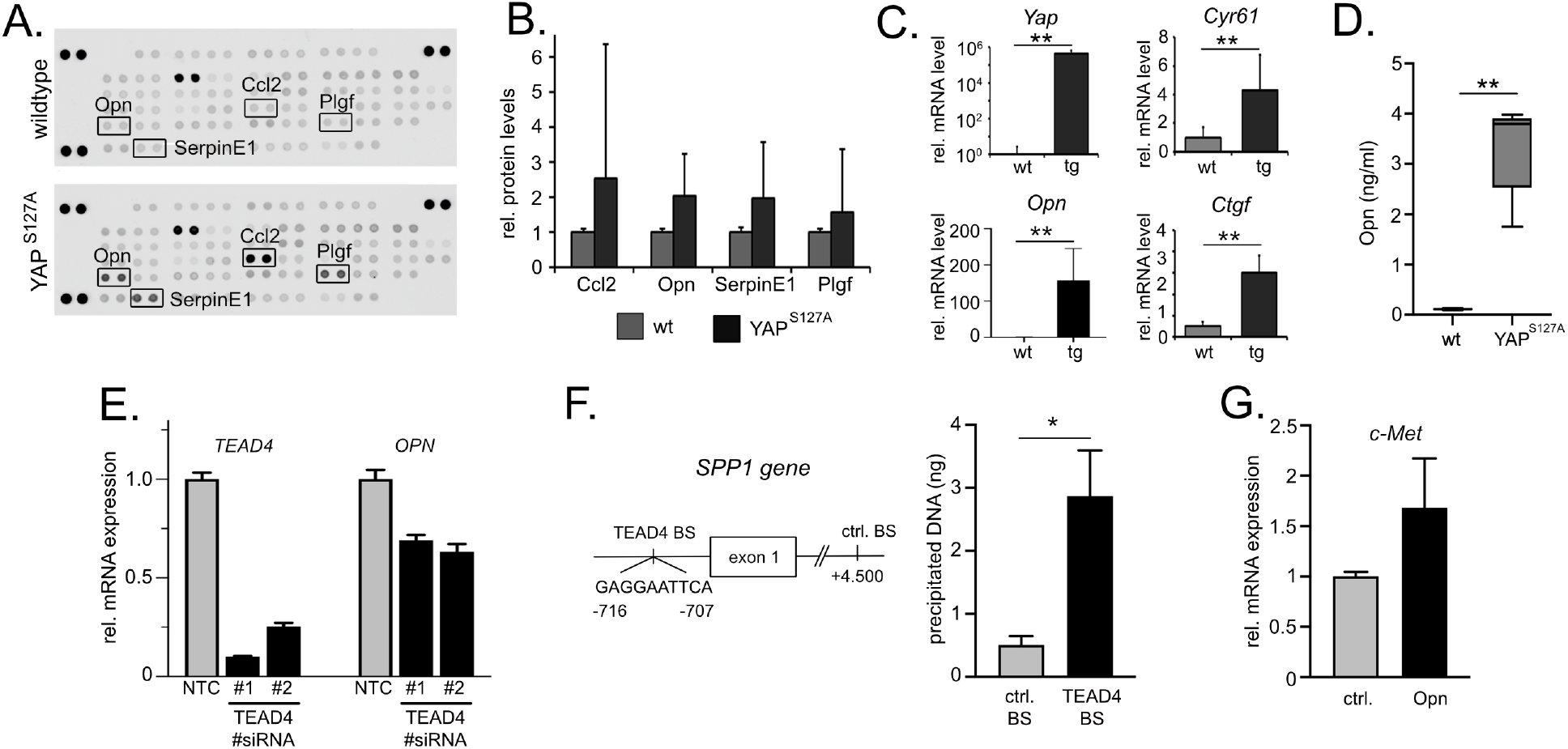
Tumor cell-derived signalling regulates heterologous EC communication. **(A.)** Exemplary proteome profiling of cell culture supernatants derived from wildtype and Yap^S127A^-transgenic hepatocytes (n=3-5/group). Candidates that were used for further analysis are indicated. **(B.)** Normalized protein levels of the four candidate factors: Ccl2, Opn, SerpinE1, Plgf (n=3-5/group). **(C.)** Confirmatory real-time PCR analysis of YAP, the positive controls Cyr61 and Ctgf as well as Opn, in samples from primary isolated hepatocytes (wt *vs*. tg), (n=5-7/group). Mann-Whitney U test (**p≤0.01). **(D.)** ELISA detecting murine Opn in mouse blood plasma derived from wildtype and Yap^S127A^-transgenic mice, (n=5-8 and n=4-5, respectively). Mann-Whitney U test, one-sided t test (**p≤0.01). **(E.)** Real-time PCR analysis of TEAD4 and OPN after siRNA-mediated silencing of TEAD4 in the liver cancer cell line Sk-Hep1. NTC: nonsense siRNA. Two independent siRNAs for TEAD4 were used (#1 and #2). One representative experiment is shown (total n=3). **(F.)** ChIP analysis of TEAD4 binding at the *OPN/SPP1* promoters at the indicated binding sites predicted by the JASPAR database. Analyses were performed with human HCC (HepG2), (n=3). Mann Whitney U test, one sided (*p≤0.05). **(G.)** OPN administration induces c-Met expression in primary liver ECs at the transcript level. One representative experiment is shown (total n=3). Student’s t-test (*p≤0.05).

For further analysis, we focused on Opn, which showed the highest dynamic range and lowest variability in the supernatants of cultured hepatocytes. Because, previous studies revealed a physical interaction between Yap and the transcription factor *TEA domain transcription factor 4* (Tead4) in hepatocytes (5), we hypothesized that Tead4 might transcriptionally regulate OPN expression through binding of gene promoter regions. Indeed, siRNA-mediated inhibition of TEAD4 reduced OPN expression in a human liver cancer cell line (Figure 5E). Furthermore, data derived from the *Encyclopedia of DNA Elements* (ENCODE) database corroborated the presence of TEAD4 binding sites within the *OPN* (syn. *SPP1*) and *CTGF* promoter (pos. control) in two cell lines (Suppl. Figure S7B). This was confirmed by chromatin immunoprecipitation (ChIP) analyses, illustrating binding of TEAD4 at the *OPN*/*SPP1* promoter predicted by a JASPAR database *in silico* screen in human HCC cells (Figure 5F) (24).

Lastly, the biological impact of Yap-dependent Opn on ECs and cells with a CECs phenotype with regard to c-Met expression was tested *in vitro* using primary isolated cells. As indicated by our previous results, OPN administration led to a moderate but consistent induction of c-Met expression in primary cells (between 23 and 69% in three independent experiments; Figure 5G).

In summary, hepatocyte-derived secreted factors such as YAP-induced Opn serve as initial effectors that induce heterologous communication networks between CECs and LSECs, for example by sensitizing CECs towards LSEC-derived Hgf.

### Endothelial changes in YAP-associated human hepatocarcinogenesis

To confirm the findings of our *in vivo* studies and screening approaches in human tissue samples, transcriptome data derived from 242 human HCC patients (HCCs and corresponding adjacent liver tissues) was analyzed (25).

A significant reduction of LSEC marker signature genes (incl. *LYVE-1* and *STAB2*) was detectable in HCCs compared to the liver tissues, while CEC markers (incl. *BCAM* and *VWF*) were elevated (Figure 6A). Interestingly, the presence of a tip cell gene signature (consisting of JAG1, LAMB1, NRP1, CXCR4, PGF, KDR, ESM1), which is characteristic of the CEC phenotype, not only significantly correlated with *OPN* transcript levels (CXCR4: r=0.34, p≤0.001; LAMB1: r=0.39, p≤0.001) but is also statistically associated with poor clinical outcome of HCC patients (Figure 3C, Figure 6B/C).

**Figure 6:**
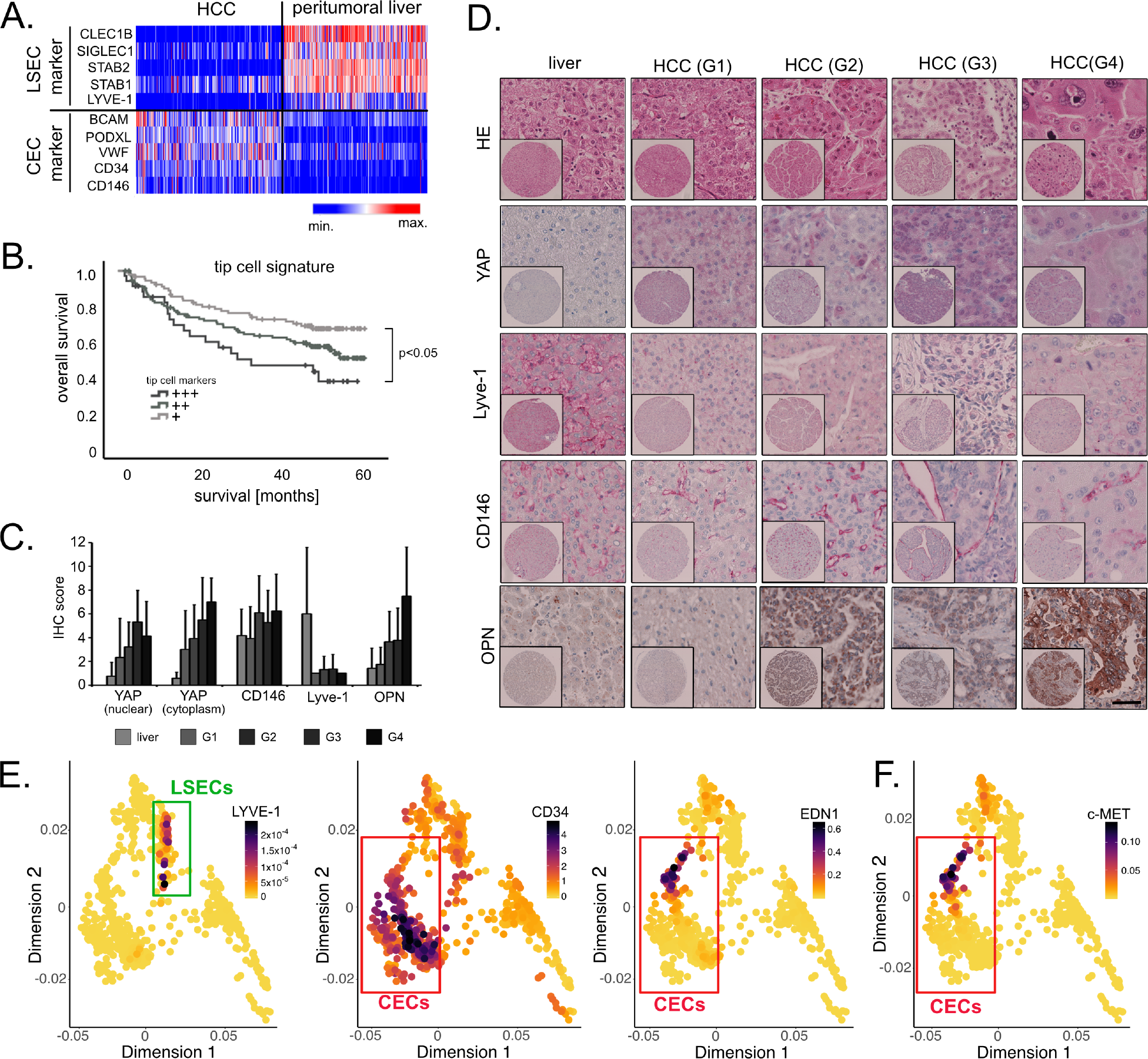
Endothelial changes in human hepatocarcinogenesis. **(A.)** Expression of 5 CEC and LSEC markers in HCCs and adjacent liver tissues. Transcriptome data derived from 242 HCC patients. **(B.)** Kaplan–Meier curves for patients with low (+), moderate (++), and high (+++) tip cell signature expression (ANOVA *p≤0.05). **(C.)** Grading dependent IHC scores of YAP (nuclear/cytoplasmic), CD146, Lyve-1, and OPN in livers and HCCs (G1-G4). **(D.)** Immunohistochemistry of YAP, its target genes OPN and the EC markers Lyve-1/CD146 using human HCC tissue microarrays (n=91 HCCs and 7 liver tissues). Exemplary liver tissue and HCCs with well (G1/G2) and poor differentiation (G3/G4) are shown. Scale bar: 50 μm. **(E.)** ScRNA-seq data derived from 12 human liver cancer samples were re-analyzed (27). Only cells that belong to the group of *tumor endothelial cells* (TECs) were regarded. Exemplary PHATE visualizations are shown with LYVE-1, CD34, and EDN1 highlighted. **(F.)** PHATE visualization for the HGF receptor c-MET is shown in a subpopulation of CD34-positive cells.

Immunohistochemical stains of liver tissues and HCCs with good (G1/G2) and poor (G3/G4) differentiation were performed (Figure 6C/D). Similar to our previous results, semiquantitative analysis revealed a progressive and statistically significant induction of YAP in cytoplasm and nuclei in hepatocarcinogenesis (r_s_=0.3; p≤0.01) (5,26). While CD146 expression only moderately increased, Lyve-1 was strongly reduced in ECs in well-differentiated HCCs (Lyve-1: r_s_=−0.26, p≤0.01). Similarly, elevated OPN abundance significantly correlated with tumor progression (OPN: r_s_=0.41; p≤0.01).

Lastly, we aimed to confirm our findings using scRNA-seq data derived from human HCC patients (27). In the group of ECs, two distinct population expressing LSEC markers (e.g. Lyve-1) and CEC markers (e.g. CD34) were identified (Figure 6E). The number of viable CEC cells was higher than of LSECs confirming the process of capillarization on these samples. Importantly, c-MET positivity was exclusively detected in ECs with CEC marker gene expression (especially EDN1) and not with LSEC markers substantiating the receptor induction in cells of the continuous endothelium (Figure 6F). In addition, in tumor cells a significant correlation between OPN and typical YAP target genes was detectable supporting the YAP-dependent regulation of OPN (e.g. CYR61: r_P_=0.49; UHMK1: r_P_=0.42; Jag-1: r_P_=0.52).

Together, these results confirmed the association between YAP activation, OPN overexpression, and the replacement of LSECs by CECs in human hepatocarcinogenesis.

## Discussion

One central goal of our study was the molecular characterization of CECs and LSECs in different stages of YAP-induced hepatocarcinogenesis. To address this, we used a model for the inducible and hepatocyte-specific expression of constitutively active Yap^S127A^ (2). This model leads to the development of hepatomegaly and rapid tumor formation within a few weeks without extensive fibrosis. Importantly, our model of Yap-induced changes in the microenvironment through a heterologous communication analysis of hepatocytes/HCC cells with ECs is supported by recent findings illustrating tumor cell-extrinsic functions of this oncogene (7). In addition, the central role of the Hippo/Yap pathway in the context of angiogenesis is discussed as the major Yap target genes Ctgf and Cyr61 are important regulators of angiogenesis (28). Moreover, Yap contributes to angiogenesis in cholangiocellular carcinoma *via* the transcriptional regulation of proangiogenic microfibrillar-associated protein 5 (MFAP5) (29).

Heterotypic communication regulates cell-type-specific functions and is required for organ development and tissue homeostasis. The liver vascular niche is a key regulator of organ functionality under physiological and pathological conditions. For example, a dynamic interlineage cross-talk in three-dimensional liver bud organoids consisting of hepatic endodermal, endothelial, and mesenchymal cells is crucial for liver development (30). EC-derived angiocrine signaling regulates organ patterning in liver development, fibrosis, and organ reconstitution after liver damage (31–33). Recent findings illustrate the relevance of ECs as integrators of mechanical forces (blood perfusion) into angiocrine signals and biological responses, such as hepatocellular proliferation and organ growth (34). Under pathophysiological conditions, the importance of ECs in tumor formation and progression has been demonstrated for various tumor entities (35); however, the impact of dynamically expressed and liver EC-derived angiokines in different stages of hepatocarcinogenesis is poorly understood.

The concept of EC heterogeneity across organs as well as their distinct molecular characteristics in homeostasis, regenerative processes, and disease is well established (36). However, little is known about the molecular characteristics of EC subtypes within tissue-specific vascular beds. For the liver, ultrastructural and histo-morphological data illustrates the existence of heterogeneous EC compartments with specific morphological features: the porto-central axis lining liver sinusoid (LSECs) and the portal/pericentral micro-vessels composed of CECs (8,9). First attempts to characterize hepatic ECs on a molecular level were conducted by comparative expression profiling of liver and lung microvascular ECs (18). This analysis revealed a gene set representative for ECs forming discontinuous micro-vessels, including transcriptional regulators (e.g. Tfec, Maf), scavenger receptors (e.g. Stabilin−1/−2) and cytokine/growth factor signaling (e.g. the WNT pathway). Recently unpaired and paired single cell analyses have resolved liver EC heterogeneity (10,19), however, a combined and detailed spatial, molecular, biochemical, and functional description of hepatic EC subtypes is not yet available. Our results demonstrate exclusive features of distinct EC populations in the liver, including the expression of signaling pathway constituents (e.g. Hippo and TGFβ pathway) and functionality (e.g. ECM interaction and cell adhesion).

Data of this study illustrate a replacement of LSECs by CECs in hepatocarcinogenesis and novel CEC subtype biomarkers including Bcam and Thbd. In combination with arterio-venous discriminating molecules such as EfnB2-EphB4 these factors may represent valuable tools for the further dissection of hepatic vascular compartments (37). This is supported by the fact publicly available single cell data derived from human HCC patients strongly suggests a further subclassification of CECs in tumor development as two CEC subtypes with distinct marker genes were detected in HCCs at the single-cell level (CD34^+^/LYVE-1^−^and EDN1^+^/LYVE-1^−^cells) (19,27). Importantly, only the EDN1^+^/LYVE-1^−^population expresses c-MET and therefore is responsive towards LSEC-derived HGF.

Hepatic vascular remodeling or ‘capillarization’ is defined by the loss of LSEC characteristics (incl. marker expression and fenestration), an increase in CEC marker expression and altered ECM deposition in the perisinusoidal space (15). Our data illustrates that this progressive shift in EC composition is detectable in the earliest phases of liver tumorigenesis. This is confirmed by previous findings showing that the loss of LSEC characteristics might be a tumor driver already in early liver disease (38). Our results also demonstrate that CEC expansion is based on active migration; however, additional cellular mechanisms may contribute to the observed EC class switch. For example, EC plasticity and endothelial-to-mesenchymal transition (EndMT) could give rise to EC populations to form a continuous endothelium (39). In addition, bone marrow-derived CSF1R^+^ cells have been identified as endothelial progenitors that integrate into existing blood vessels (40,41).

The dynamic and tumor stage-dependent expression patterns derived from CECs and LSECs led us draw several conclusions. Firstly, Yap induces a rapid heterologous communication network, which is already detectable in early and pre-malignant stages of hepatocarcinogenesis. This network is central for remodeling the vascular niche, which may be a mandatory step for the ‘angiogenic switch’ needed for tumor progression (42). Secondly, angiocrine signaling not only connects ECs with parenchymal hepatocytes (33) or non-parenchymal HSCs (43), but may also form a communication network between EC subtypes. Interestingly, the dynamic regulation of ligand/receptor pairs in CECs and LSECs is suggestive of a multifactorial connection between CECs and LSECs (e.g. Hgf/c-Met, Vegfc/Vegfr3, Cxcl12/Cxcr4), which may affect their spatio-temporal responsiveness and their biological properties. Different studies have already illustrated that LSECs represent the major cellular source for Hgf (31,34). However, the progressive increase of c-Met in CECs, in combination with constitutively high HGF production by LSECs, raised our special interest, as it indicated that CECs gain their responsiveness towards Hgf in liver cancer development. Indeed, the relevance of the Hgf/c-Met axis in aberrant vascularization e.g. in neural tumor development and during liver regeneration has been confirmed in previous work (31,44). It is worth mentioning that ECM constituents with low-affinity binding sites for Hgf (e.g. collagens) may serve as source for this growth factor (45). Thus, LSEC-derived Hgf could provide an important origin for ECM-bound Hgf, which is released upon request in the course of capillarization in hepatocarcinogenesis.

Induction of the YAP-dependent heterologous communication network in our mouse model must originate from genetically-modified hepatocytes. Among the hepatocyte-derived factors, Opn is transcriptionally regulated by Yap/Tead4 and induces c-Met expression in liver ECs. Based on our results, we conclude that Opn represents one communication hub between tumor cells and CECs, which sensitizes this EC subtype for LSEC-derived Hgf. Indeed, the relevance of Opn as a central regulator of angiogenesis has already been confirmed by several studies, such as in pancreatic cancer and breast cancer (46,47). If YAP-dependent extrinsic functions control heterologous communication and capillarization in other HCC models that equally show YAP overexpression (e.g. AlbTag and Hep56.1D), or whether YAP-independent mechanisms contribute to this phenotype, needs to be clarified in future studies.

## Abbreviations

CEC: capillary endothelial cell
EC: endothelial cell
ECM: extracellular matrix
HCC: hepatocellular carcinoma
Hgf/HGF^*^: hepatocyte growth factor
HSC: hepatic stellate cell
LSEC: liver sinusoidal endothelial cell
NPC: non-parenchymal cell
Opn: osteopontin
Yap/YAP^*^: yes-associated protein

## Acknowledgement

The authors thank Jennifer Schmitt, Michaela Bissinger, Sandra Förmer, Jutta Scheuerer, and Nina Hofmann for excellent technical assistance. We also want to thank F. D. Camargo for providing the Col1A1-Yap^S127A^ transgenic mice. Tissue samples were provided by the Tissue Bank of the National Center of Tumor Diseases (NCT, Heidelberg, Germany) in accordance with the regulations of the Tissue Bank and the approval of the Ethics Committee of Heidelberg University. We thank the Center of Model System and Comparative Pathology (CMCP; Tanja Poth) and Heike Conrad, Diana Lutz, Karin Rebholz, and Veronica Geissler for their support. We also want to thank Ulrike Ganserer and Dr. Ingrid Hausser for conducting scanning electron microscopy as well as Martin Dittmer for providing us with support in programming. We want to thank Katherine Zach for manuscript proof reading.

## Footnotes

* upper and lower cases were used according to the murine or human origin

## References

1. Patel SH, Camargo FD, Yimlamai D. Hippo Signaling in the Liver Regulates Organ Size, Cell Fate, and Carcinogenesis. Gastroenterology 2017;152:533–45

2. Camargo FD, Gokhale S, Johnnidis JB, Fu D, Bell GW, Jaenisch R, et al. YAP1 increases organ size and expands undifferentiated progenitor cells. Curr Biol 2007;17:2054–60

3. Yimlamai D, Christodoulou C, Galli GG, Yanger K, Pepe-Mooney B, Gurung B, et al. Hippo pathway activity influences liver cell fate. Cell 2014;157:1324–38

4. Cabochette P, Vega-Lopez G, Bitard J, Parain K, Chemouny R, Masson C, et al. YAP controls retinal stem cell DNA replication timing and genomic stability. Elife 2015;4:e08488

5. Weiler SME, Pinna F, Wolf T, Lutz T, Geldiyev A, Sticht C, et al. Induction of Chromosome Instability by Activation of Yes-Associated Protein and Forkhead Box M1 in Liver Cancer. Gastroenterology 2017;152:2037–51 e22

6. Yang S, Zhang L, Liu M, Chong R, Ding SJ, Chen Y, et al. CDK1 phosphorylation of YAP promotes mitotic defects and cell motility and is essential for neoplastic transformation. Cancer Res 2013;73:6722–33

7. Guo X, Zhao Y, Yan H, Yang Y, Shen S, Dai X, et al. Single tumor-initiating cells evade immune clearance by recruiting type II macrophages. Genes Dev 2017;31:247–59

8. Potente M, Makinen T. Vascular heterogeneity and specialization in development and disease. Nat Rev Mol Cell Biol 2017;18:477–94

9. Aird WC. Phenotypic heterogeneity of the endothelium: II. Representative vascular beds. Circ Res 2007;100:174–90

10. Halpern KB, Shenhav R, Massalha H, Toth B, Egozi A, Massasa EE, et al. Paired-cell sequencing enables spatial gene expression mapping of liver endothelial cells. Nat Biotechnol 2018;36:962–70

11. Limmer A, Ohl J, Kurts C, Ljunggren HG, Reiss Y, Groettrup M, et al. Efficient presentation of exogenous antigen by liver endothelial cells to CD8+ T cells results in antigen-specific T-cell tolerance. Nat Med 2000;6:1348–54

12. Strauss O, Phillips A, Ruggiero K, Bartlett A, Dunbar PR. Immunofluorescence identifies distinct subsets of endothelial cells in the human liver. Sci Rep 2017;7:44356

13. Rocha AS, Vidal V, Mertz M, Kendall TJ, Charlet A, Okamoto H, et al. The Angiocrine Factor Rspondin3 Is a Key Determinant of Liver Zonation. Cell Rep 2015;13:1757–64

14. Geraud C, Evdokimov K, Straub BK, Peitsch WK, Demory A, Dorflinger Y, et al. Unique cell type-specific junctional complexes in vascular endothelium of human and rat liver sinusoids. PLoS One 2012;7:e34206

15. Geraud C, Mogler C, Runge A, Evdokimov K, Lu S, Schledzewski K, et al. Endothelial transdifferentiation in hepatocellular carcinoma: loss of Stabilin-2 expression in peritumourous liver correlates with increased survival. Liver Int 2013;33:1428–40

16. DeLeve LD. Liver sinusoidal endothelial cells in hepatic fibrosis. Hepatology 2015;61:1740–6

17. Xie G, Choi SS, Syn WK, Michelotti GA, Swiderska M, Karaca G, et al. Hedgehog signalling regulates liver sinusoidal endothelial cell capillarisation. Gut 2013;62:299–309

18. Geraud C, Schledzewski K, Demory A, Klein D, Kaus M, Peyre F, et al. Liver sinusoidal endothelium: a microenvironment-dependent differentiation program in rat including the novel junctional protein liver endothelial differentiation-associated protein-1. Hepatology 2010;52:313–26

19. Aizarani N, Saviano A, Sagar, Mailly L, Durand S, Herman JS, et al. A human liver cell atlas reveals heterogeneity and epithelial progenitors. Nature 2019;572:199–204

20. del Toro R, Prahst C, Mathivet T, Siegfried G, Kaminker JS, Larrivee B, et al. Identification and functional analysis of endothelial tip cell-enriched genes. Blood 2010;116:4025–33

21. Rocha SF, Schiller M, Jing D, Li H, Butz S, Vestweber D, et al. Esm1 modulates endothelial tip cell behavior and vascular permeability by enhancing VEGF bioavailability. Circ Res 2014;115:581–90

22. Strasser GA, Kaminker JS, Tessier-Lavigne M. Microarray analysis of retinal endothelial tip cells identifies CXCR4 as a mediator of tip cell morphology and branching. Blood 2010;115:5102–10

23. O’Connell KA, Edidin M. A mouse lymphoid endothelial cell line immortalized by simian virus 40 binds lymphocytes and retains functional characteristics of normal endothelial cells. J Immunol 1990;144:521–5

24. Mathelier A, Zhao X, Zhang AW, Parcy F, Worsley-Hunt R, Arenillas DJ, et al. JASPAR 2014: an extensively expanded and updated open-access database of transcription factor binding profiles. Nucleic Acids Res 2014;42:D142–7

25. Roessler S, Jia HL, Budhu A, Forgues M, Ye QH, Lee JS, et al. A unique metastasis gene signature enables prediction of tumor relapse in early-stage hepatocellular carcinoma patients. Cancer Res 2010;70:10202–12

26. Tschaharganeh DF, Chen X, Latzko P, Malz M, Gaida MM, Felix K, et al. Yes-associated protein up-regulates Jagged-1 and activates the Notch pathway in human hepatocellular carcinoma. Gastroenterology 2013;144:1530–42 e12

27. Ma L, Hernandez MO, Zhao Y, Mehta M, Tran B, Kelly M, et al. Tumor Cell Biodiversity Drives Microenvironmental Reprogramming in Liver Cancer. Cancer Cell 2019;36:418–30 e6

28. Brigstock DR. Regulation of angiogenesis and endothelial cell function by connective tissue growth factor (CTGF) and cysteine-rich 61 (CYR61). Angiogenesis 2002;5:153–65

29. Marti P, Stein C, Blumer T, Abraham Y, Dill MT, Pikiolek M, et al. YAP promotes proliferation, chemoresistance, and angiogenesis in human cholangiocarcinoma through TEAD transcription factors. Hepatology 2015;62:1497–510

30. Camp JG, Sekine K, Gerber T, Loeffler-Wirth H, Binder H, Gac M, et al. Multilineage communication regulates human liver bud development from pluripotency. Nature 2017;546:533–8

31. Ding BS, Nolan DJ, Butler JM, James D, Babazadeh AO, Rosenwaks Z, et al. Inductive angiocrine signals from sinusoidal endothelium are required for liver regeneration. Nature 2010;468:310–5

32. Geraud C, Koch PS, Zierow J, Klapproth K, Busch K, Olsavszky V, et al. GATA4-dependent organ-specific endothelial differentiation controls liver development and embryonic hematopoiesis. J Clin Invest 2017;127:1099–114

33. Hu J, Srivastava K, Wieland M, Runge A, Mogler C, Besemfelder E, et al. Endothelial cell-derived angiopoietin-2 controls liver regeneration as a spatiotemporal rheostat. Science 2014;343:416–9

34. Lorenz L, Axnick J, Buschmann T, Henning C, Urner S, Fang S, et al. Mechanosensing by beta1 integrin induces angiocrine signals for liver growth and survival. Nature 2018;562:128–32

35. Wieland E, Rodriguez-Vita J, Liebler SS, Mogler C, Moll I, Herberich SE, et al. Endothelial Notch1 Activity Facilitates Metastasis. Cancer Cell 2017;31:355–67

36. Nolan DJ, Ginsberg M, Israely E, Palikuqi B, Poulos MG, James D, et al. Molecular signatures of tissue-specific microvascular endothelial cell heterogeneity in organ maintenance and regeneration. Dev Cell 2013;26:204–19

37. Adams RH, Wilkinson GA, Weiss C, Diella F, Gale NW, Deutsch U, et al. Roles of ephrinB ligands and EphB receptors in cardiovascular development: demarcation of arterial/venous domains, vascular morphogenesis, and sprouting angiogenesis. Genes Dev 1999;13:295–306

38. Xie G, Wang X, Wang L, Wang L, Atkinson RD, Kanel GC, et al. Role of differentiation of liver sinusoidal endothelial cells in progression and regression of hepatic fibrosis in rats. Gastroenterology 2012;142:918–27 e6

39. Dejana E, Hirschi KK, Simons M. The molecular basis of endothelial cell plasticity. Nat Commun 2017;8:14361

40. Plein A, Fantin A, Denti L, Pollard JW, Ruhrberg C. Erythro-myeloid progenitors contribute endothelial cells to blood vessels. Nature 2018;562:223–8

41. Singhal M, Liu X, Inverso D, Jiang K, Dai J, He H, et al. Endothelial cell fitness dictates the source of regenerating liver vasculature. J Exp Med 2018;215:2497–508

42. Baeriswyl V, Christofori G. The angiogenic switch in carcinogenesis. Semin Cancer Biol 2009;19:329–37

43. Marrone G, Shah VH, Gracia-Sancho J. Sinusoidal communication in liver fibrosis and regeneration. J Hepatol 2016;65:608–17

44. Huang M, Liu T, Ma P, Mitteer RA, Jr., Zhang Z, Kim HJ, et al. c-Met-mediated endothelial plasticity drives aberrant vascularization and chemoresistance in glioblastoma. J Clin Invest 2016;126:1801–14

45. Schuppan D, Schmid M, Somasundaram R, Ackermann R, Ruehl M, Nakamura T, et al. Collagens in the liver extracellular matrix bind hepatocyte growth factor. Gastroenterology 1998;114:139–52

46. You WK, Sennino B, Williamson CW, Falcon B, Hashizume H, Yao LC, et al. VEGF and c-Met blockade amplify angiogenesis inhibition in pancreatic islet cancer. Cancer Res 2011;71:4758–68

47. Raja R, Kale S, Thorat D, Soundararajan G, Lohite K, Mane A, et al. Hypoxia-driven osteopontin contributes to breast tumor growth through modulation of HIF1alpha-mediated VEGF-dependent angiogenesis. Oncogene 2014;33:2053–64

