## supplementary material for "YAP orchestrates heterotypic endothelial cell communication via HGF/c-MET signaling in liver tumorigenesis"

*S. Thomann et al., - suppl. figure S1*

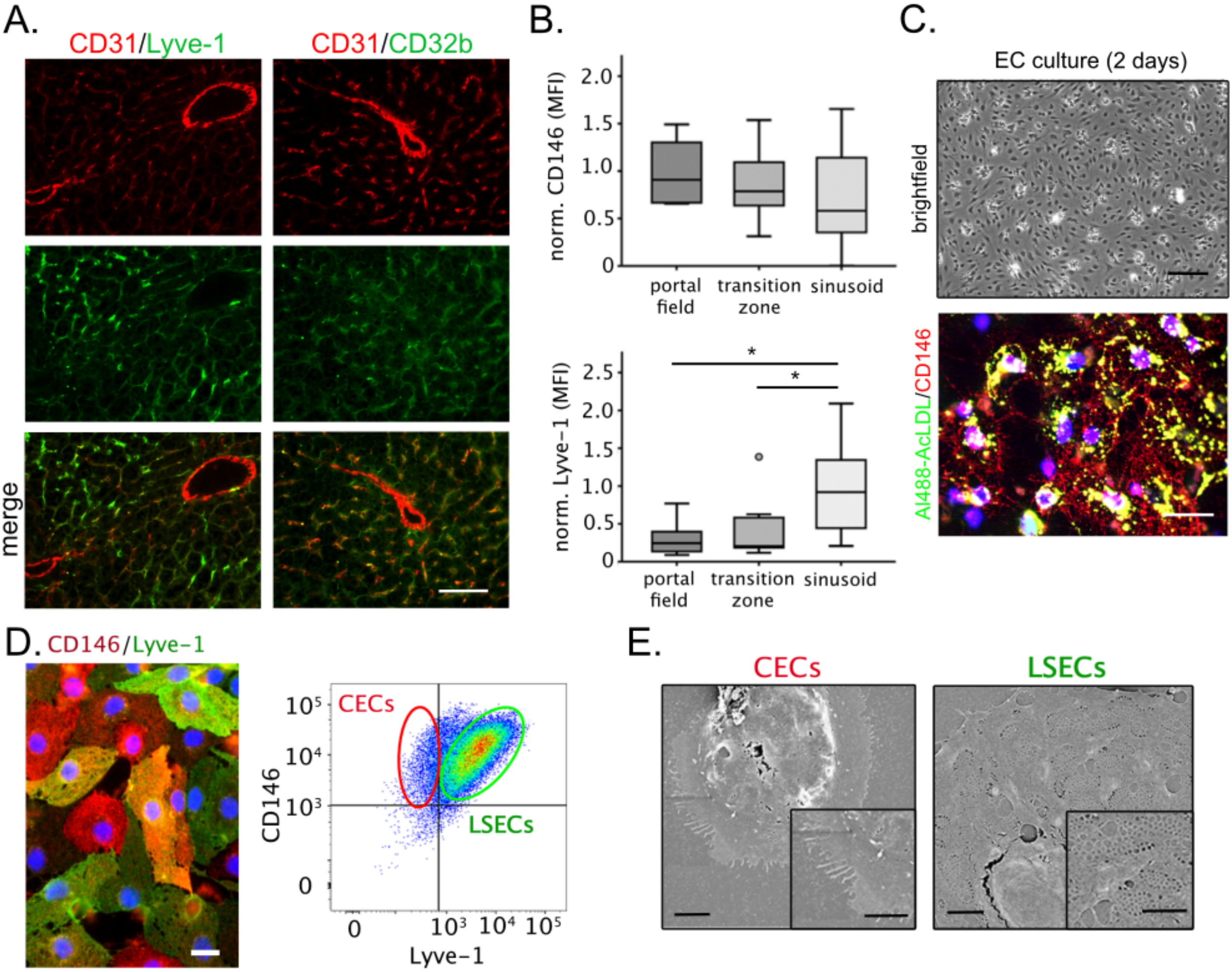

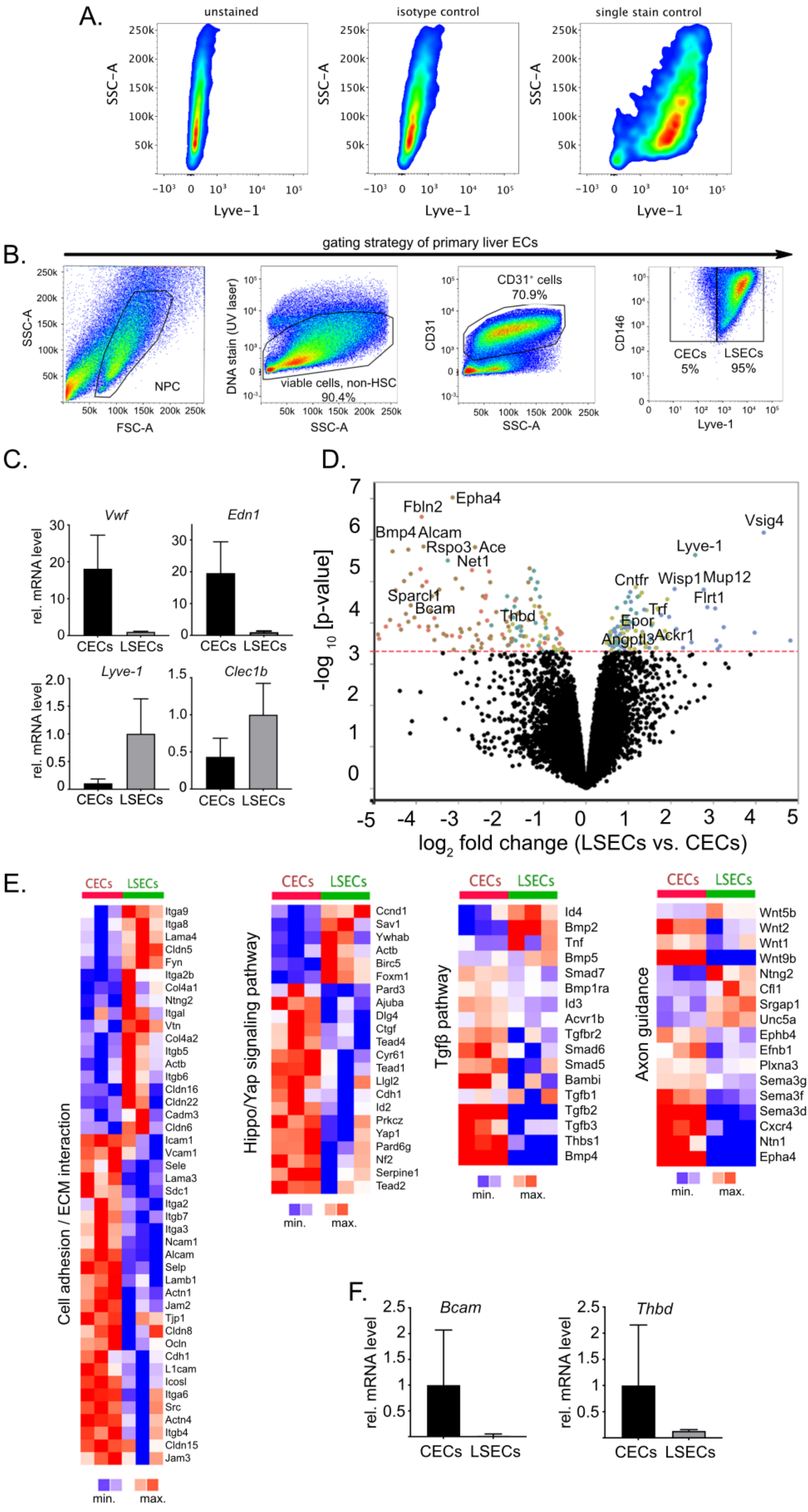
A.
unstained
isotype control
single stain control
SSC-A
Lyve-1
B.
gating strategy of primary liver ECs
NPC
viable cells, non-HSC
90.4%
CD31⁺ cells
70.9%
CECs
5%
LSECs
95%
FSC-A
SSC-A
DNA stain (UV laser)
CD31
CD146
Lyve-1
C.
Vwf
Edn1
Lyve-1
Clec1b
rel. mRNA level
CECs LSECs
D.
-log10 [p-value]
log2 fold change (LSECs vs. CECs)
Fbln2
Epha4
Bmp4
Alcam
Rspo3
Ace
Net1
Sparcl1
Bcam
Thbd
Vsig4
Lyve-1
Cntfr
Wisp1
Mup12
Flrt1
Trf
Epor
Angptl3
Ackr1
E.
CECs LSECs
Cell adhesion / ECM interaction
Itga9 Itga8 Lama4 Cldn5 Fyn Itga2b Col4a1 Ntng2 Itgal Vtn Col4a2 Itgb5 Actb Itgb6 Cldn16 Cldn22 Cadm3 Cldn6 Icam1 Vcam1 Sele Lama3 Sdc1 Itga2 Itgb7 Itga3 Ncam1 Alcam Selp Lamb1 Actn1 Jam2 Tjp1 Cldn8 Ocln Cdh1 L1cam Icosl Itga6 Src Actn4 Itgb4 Cldn15 Jam3
Hippo/Yap signaling pathway
Ccnd1 Sav1 Ywhab Actb Birc5 Foxm1 Pard3 Ajuba Dlg4 Ctgf Tead4 Cyr61 Tead1 Llgl2 Cdh1 Id2 Prkcz Yap1 Pard6g Nf2 Serpine1 Tead2
Tgfβ pathway
Id4 Bmp2 Tnf Bmp5 Smad7 Bmp1ra Id3 Acvr1b Tgfbr2 Smad6 Smad5 Bambi Tgfb1 Tgfb2 Tgfb3 Thbs1 Bmp4
Axon guidance
Wnt5b Wnt2 Wnt1 Wnt9b Ntng2 Cfl1 Srgap1 Unc5a Ephb4 Efnb1 Plxna3 Sema3g Sema3f Sema3d Cxcr4 Ntn1 Epha4
min. max.
F.
Bcam
Thbd
rel. mRNA level
CECs LSECs

### *S. Thomann et al., - suppl. figure S3*

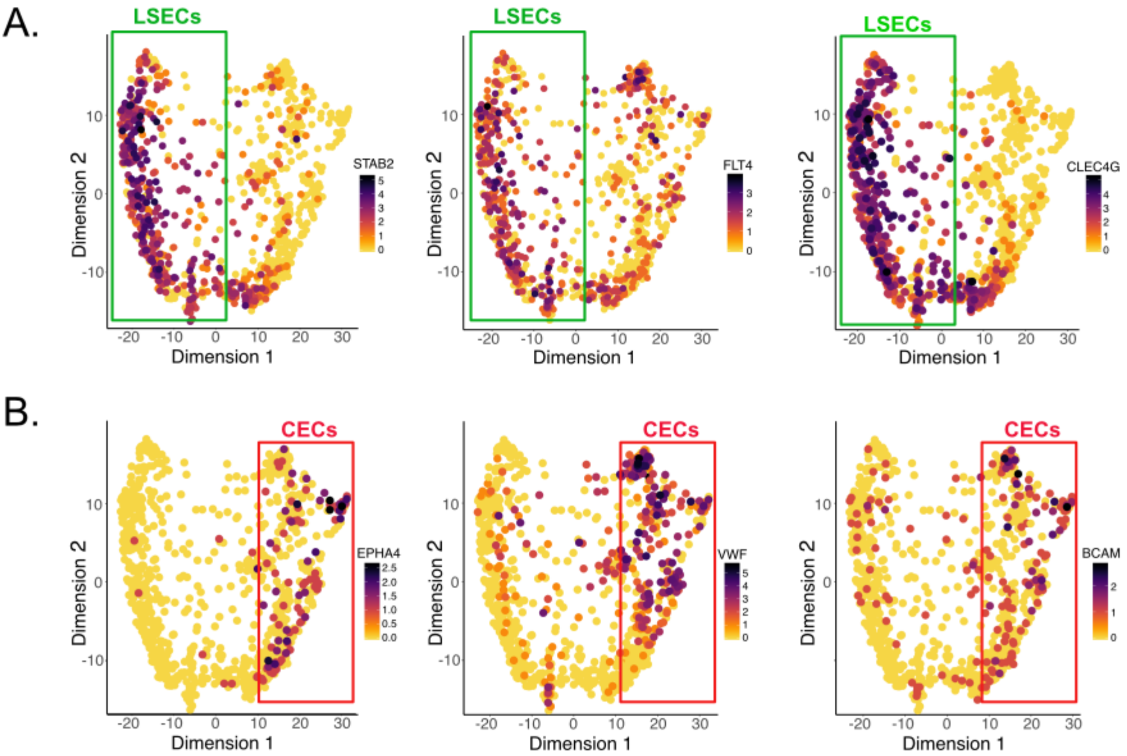

*S. Thomann et al., - suppl. figure S4*

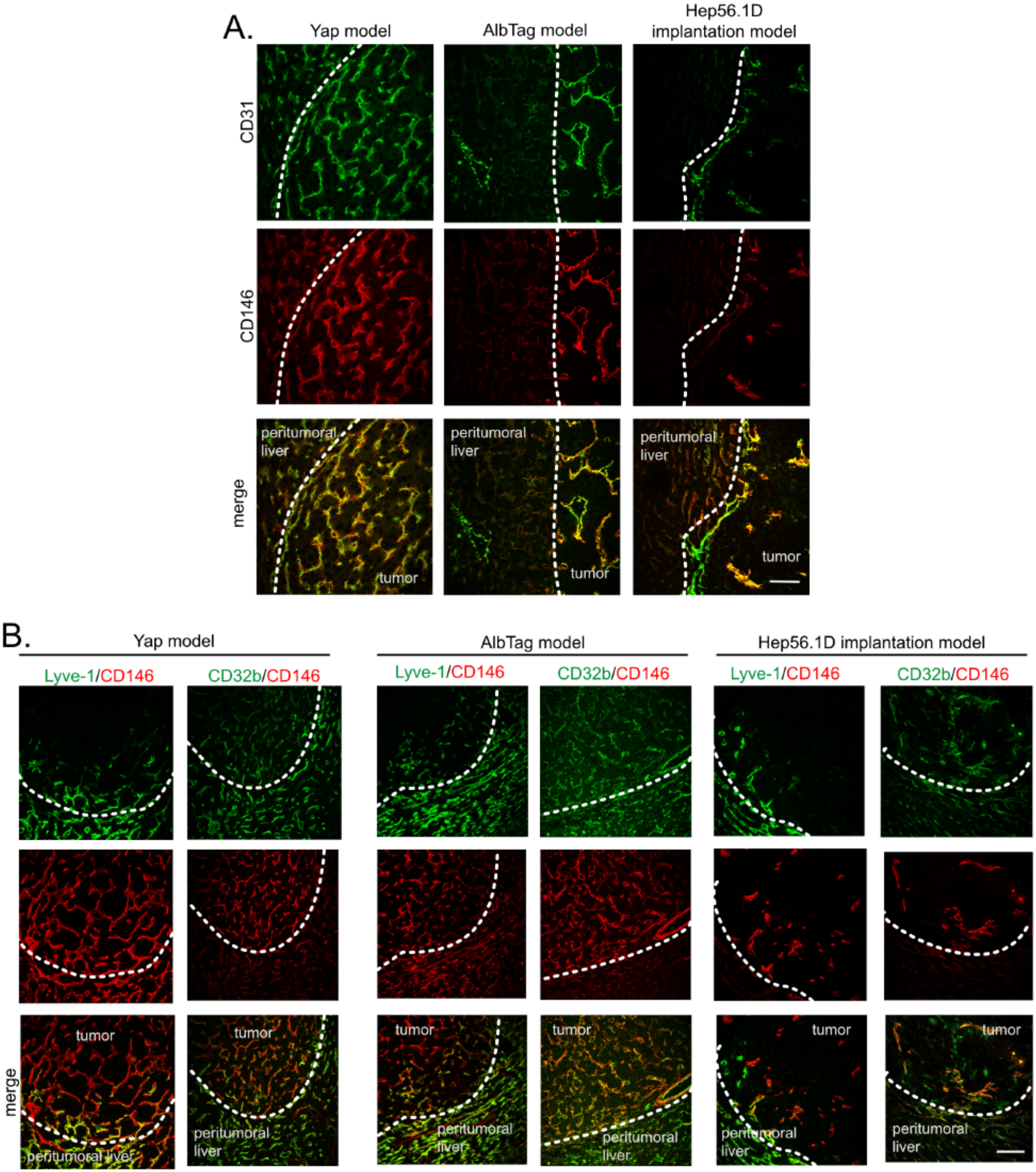

*S. Thomann et al., - suppl. figure S5*

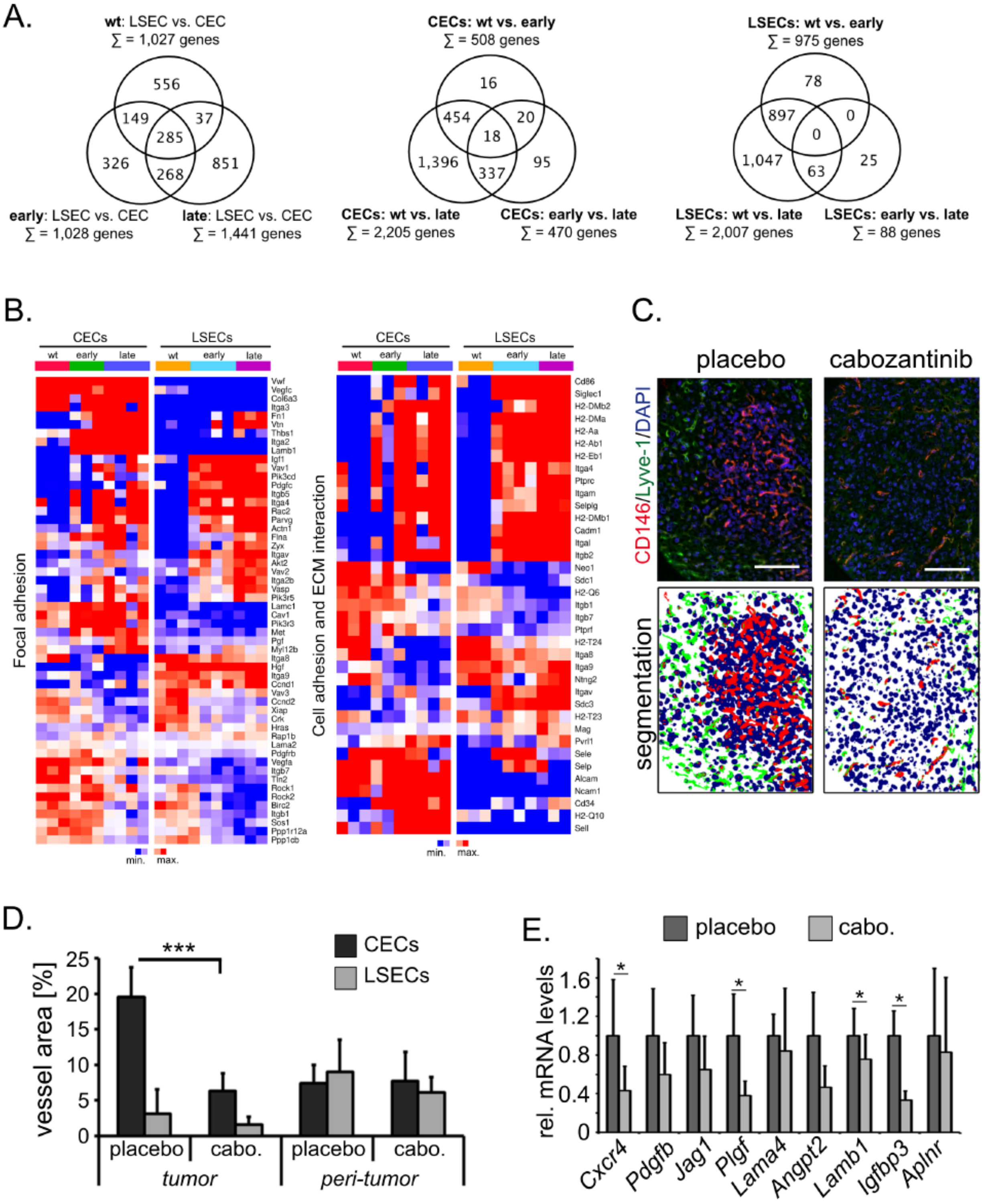

*S. Thomann et al., - suppl. figure S6*

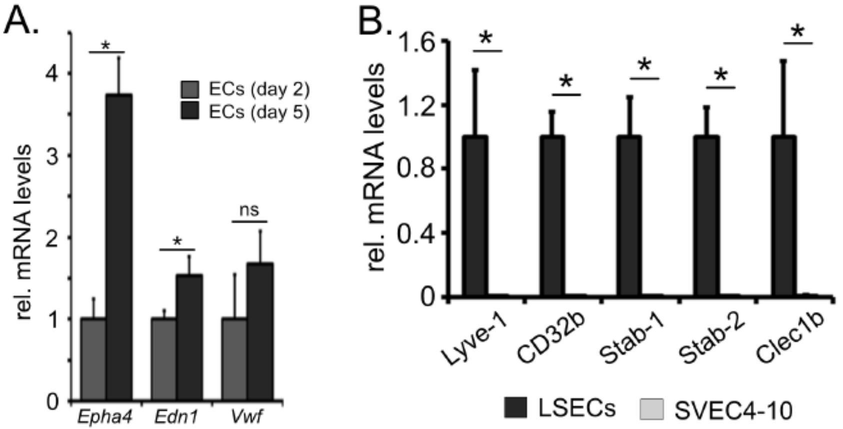


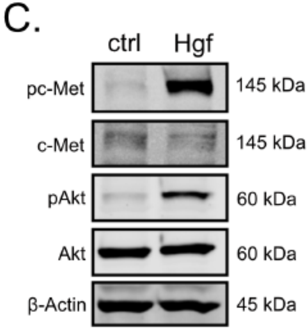


D.

| functional groups | gene count | NES |
|---|---|---|
| angiogenesis | 48 | 2.5 |
| positive regulation of cell migration | 41 | 2.5 |
| heart development | 40 | 1.9 |
| nervous system development | 40 | 1.3 |
| cell migration | 36 | 2.3 |
| positive regulation of angiogenesis | 21 | 2.1 |
| wound healing | 19 | 2.5 |
| regulation of cell shape | 19 | 1.7 |
| axon guidance | 18 | 1.5 |
| blood vessel development | 12 | 2.1 |
| positive regulation of cell adhesion | 11 | 2.4 |
| response to wounding | 11 | 2.3 |
| microtubule-based movement | 11 | 1.7 |
| blood vessel morphogenesis | 10 | 3.5 |

S. Thomann et al., - suppl. figure S7

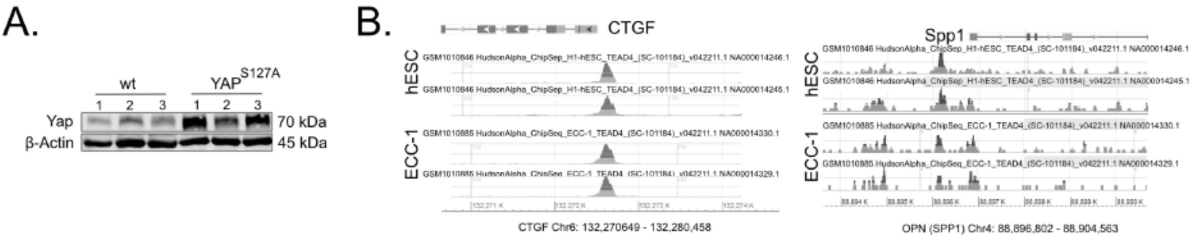

### YAP orchestrates heterotypic endothelial cell communication via HGF/c-MET signaling in liver tumorigenesis

S. Thomann, S.M.E. Weiler, S. Marquard, C.R. Ball, M. Tóth, T. Wei, C. Sticht, C. De La Torre, E. Ryschich, O. Ermakova, C. Mogler, D. Kazdal, N. Gretz, H. Glimm, E. Rempel, P. Schirmacher, and K. Breuhahn

**Figure Legends**

**Suppl. Figure S1: Morphological and functional characterization of hepatic EC heterogeneity**

**(A.)** Confirmatory characterization of distinct EC populations in healthy liver. CD31 (red) and Lyve-1/ CD32b (green) double IF stains characterize CECs ($CD31^{++}$/$Lyve\text{-}1^{-}$ or $CD31^{++}$/$CD32b^{-}$) and LSECs ($CD31^{+}$/$Lyve\text{-}1^{+}$ or $CD31^{+}$/$CD32b^{+}$). Scale bar: 50 µm.

**(B.)** Normalized MFI measurement of CD146 and Lyve-1 in portal fields, porto-sinusoidal transition zones, and sinusoids are depicted. Data are shown as box plots (n=10), statistical test: ANOVA (*p≤0.05).

**(C.)** Brightfield image of primary isolated ECs with typical cobblestone morphology. Scale bar: 50 µm. Uptake of Al488-labeled AcLDL (green) by primary ECs (CD146, red) was detected two days after isolation. Scale bar: 20 µm

**(D.)** Double IF stains for CD146 (red) and Lyve-1 (green) of primary ECs 48 hours after isolation (left panel). Exemplary FACS-based quantification of CECs ($CD146^{+}$, $Lyve\text{-}1^{-}$) and LSECs ($CD146^{+}$, $Lyve\text{-}1^{+}$), (right panel). Scale bar: 20 µm.

**(E.)** Representative scanning electron microscopic pictures confirm the presence of sieve plate fenestrations in most ECs (representing LSECs), while CECs lack respective fenestrations. Scale bars in lower right corner: 4 µm.

**Suppl. Figure S2: Isolation and characterization of CECs and LSECs**

**(A.)** Representative FACS labeling controls for Lyve-1 compensation and definition of gates. Unstained, isotype stained and single stained positive EC controls were used in each experiment. Identical controls exist for the markers used in this study (CD31, CD146).

**(B.)** Exemplary FACS-based isolation of highly pure CEC and LSEC fractions from a MACS-pre-purified and viable NPC population. After exclusion of non-viable and retinoid-positive cells (HSCs), an additional purification step for ECs *via* CD31 was performed. The final separation of CECs and LSECs was achieved using CD146/Lyve1. Purity is indicated in percent.

**(C.)** Analysis of subtype marker expression by real-time PCR. Higher expression of CEC markers (Vwf, Edn1; black bars) and LSEC markers (Lyve-1, Clec1b; grey bars) in the cell-populations confirm successful sorting procedure. Graphs show mean ± SD (n=4-7).

**(D.)** Volcano plot depicting 247 genes that are differentially expressed between CECs and LSECs (140 elevated genes in CECs and 107 elevated genes in LSECs). Typical LSEC- and CEC-specific denominators are indicated. p-values (y-axis) are shown as $\log_{10}$, while relative transcript changes are shown as $\log_2$ (x-axis), (n=3).

**(E.)** Heatmaps showing significantly regulated genes involved in ECM interaction, Hippo/YAP and Tgfβ signaling as well as axon guidance (KEGG pathways mmu04512/04514, mmu04390, mmu04350, mmu04360), (n=3).

**(F.)** Confirmatory real-time PCR analysis of Bcam and Thbd in primary isolated fractions of CECs and LSECs. Graphs show mean ± SD (n=3).

**Suppl. Figure S3: Analysis of scRNA-seq data derived from healthy liver tissues**

**(A.)** PHATE visualization of LSEC marker genes STAB2, FLT4, and CLEC4G in all ECs derived from 9 liver tissues.

**(B.)** PHATE visualization of CEC marker genes EPHA4, VWF, and BCAM in all ECs.

**Suppl. Figure S4: Dynamic EC subpopulation changes in liver tumor models**

**(A.)** Confocal CD31 (green) and CD146 (red) IF stains of three mouse liver tumor models (autochthonous $Yap^{S127A}$, AlbTag, and orthotopic Hep56.1D implantation). Tumor vessels are characterized by strong CD146 and CD31 positivity. Scale bar: 50 µm, n=3-4 animals/group.

**(B.)** CD146 (red) and Lyve-1 (green) as well as CD146 (red) and CD32b (green) IF stains in tumor tissues derived from $Yap^{S127A}$, AlbTag, and Hep56.1D implantation mouse models. Loss of LSEC-specific Lyve-1 and CD32b expression is detected in tumor tissues. Scale bar: 50 µm, n=3-4/group.

**Suppl. Figure S5: Molecular characterization of LSECs and CECs in YAP-induced tumorigenesis**

**(A.)** Venn diagrams illustrating the differential gene expression between CECs and LSECs at different stages of oncogene-induced tumorigenesis or between different liver tissues: normal (wt), hyperplastic (early), and tumor-bearing (late) livers. n=3-4/group.

**(B.)** Hierarchically clustered gene expression data of the KEGG pathway constituents 'focal adhesion – mmu04510' and 'cell adhesion and ECM interaction – mmu04512/04514)'; n=3-4/group.

**(C.)** Cabozantinib was injected for two weeks in $Yap^{S127A}$-transgenic mice. CD146/Lyve-1 IF followed by an intensity-based segmentation revealed a significant reduction of CECs but not LSEC vessel areas within tumor nodules, whereas CEC and LSEC abundance was not significantly affected in peri-tumorous liver. Channel-based segmentation of Lyve-1 (green), CD146 (red), and DAPI (blue) stained liver tissues derived from $Yap^{S127A}$-transgenic mice after placebo and cabozantinib treatment. Scale bar: 200 µm.

**(D.)** Image channel-based segmentation and vessel area calculation of intra-tumoral and peri-tumoral Lyve-1 and CD146 positive areas. Graphs show mean ± SD (10 images/group). Mann-Whitney U test (***p≤0.001).

**(E.)** Relative transcript levels of 9 tip cell markers in total livers samples after cabozantinib or placebo treatment. Graphs show mean ± SD (n=6/group); Mann-Whitey U test (*p≤0.05).

**Suppl. Figure S6: HGF/c-MET signaling in a CEC model system**

**(A.)** Real time PCR of primary isolated ECs cultured for 2 (grey) and 5 days (dark grey). Expression of the CEC markers Epha4, Edn1, and Vwf was analyzed. Graphs show mean ± SD (n=4); Mann-Whitney-U test (*p≤0.05, ns - not significant).

**(B.)** Real-time PCR-based comparison of known LSEC markers in primary LSECs and immortalized

SVEC4-10 cells. Graphs show mean ± SD (n=4), Mann-Whitney U test (*$p \leq 0.05$).

**(C.)** Western immunoblot detecting total AKT and c-Met as well as respective phospho-proteins (Ser473 for pAKT and Tyr1234/1235 pc-Met) in serum-starved SVEC4-10 cells 20 minutes after HGF administration (20 ng/ml). Actin was used as loading control.

**(D.)** Significantly upregulated genes 3 and 6 hours after HGF administration in SVEC4-10 cells were used for functional annotation analysis (DAVID). Table shows functional groups (GO-ID; biological process) according to the gene number and the corresponding normalized enrichments scores (NES).

**Suppl. Figure S7: Tumor cell-derived soluble factors are transcriptionally regulated by TEAD**

**(A.)** Western immunoblotting of Yap protein levels derived from primary isolated hepatocytes (wt and $Yap^{S127A}$). β-actin was detected for sample normalization, n=3/group.

**(B.)** TEAD4-ChIP sequencing data of the ENCODE database for the YAP target genes OPN in the human cell lines hESC and ECC1 (GSM1010845 and GSM1010885). CTGF data is included as positive control.

### YAP orchestrates heterotypic endothelial cell communication via HGF/c-MET signaling in liver tumorigenesis

S. Thomann, S.M.E. Weiler, S. Marquard, C.R. Ball, M. Tóth, T. Wei, C. Sticht, C. De La Torre, E. Ryschich, O. Ermakova, C. Mogler, D. Kazdal, N. Gretz, H. Glimm, E. Rempel, P. Schirmacher, and K. Breuhahn

#### Supplemental Material and Methods

##### Key Resources Table

| Primary Antibody (clone) | Source/Company | Identifier |
|---|---|---|
| Actb (C4) | MP Biomedicals (Santa Ana, USA) | ICN691001 |
| AKT | Cell Signaling (Cambridge, UK) | #9272; RRID: AB_329827 |
| pAKT (D9E) | Cell Signaling | #4060; RRID: AB_2315049 |
| Bcam | R&D Systems (Minneapolis, USA) | AF8299 |
| CD11b (m1/70.1+A6:C285.11.5) | Miltenyi Biotec (Bergisch Gladbach) | #130-091-241; RRID: AB_244268 |
| CD11b (EPR1344) | Abcam | ab133357; RRID: AB_2650514 |
| CD31 (390) | Biolegend (San Diego, USA) | #102502; RRID: AB_2566676 |
| CD31 | Dako (Jena, Germany) | M0823; RRID: AB_2114471 |
| CD32b (190907) | R&D Systems | MAB14601; RRID: AB_2103729 |
| CD146 (ME-9F1) | Biolegend | #134702; RRID: AB_2721501 |
| CD146 | Atlas Antibodies | HPA008848; RRID: AB_1078445 |
| Lyve-1 (Aly7) | eBioscience | #14-0443-82; RRID: AB_1724157 |
| c-Met (25H2) | Cell Signaling | #3127; RRID: AB_331361 |
| pc-Met (3D7) | Cell Signaling | #3129; RRID: AB_561173 |
| TEAD4 (N-G2) | Santa Cruz (Dallas, USA) | sc-101184; RRID: AB_2203086 |
| Thbd (461714) | R&D Systems | MAB3894; RRID: AB_920518 |
| YAP (D8H1X) | Cell Signaling | #14074; RRID: AB_2650491 |
| YAP | Cell Signaling | #4912; RRID: AB_2218911 |

| Secondary Antibody | Source/Company | Identifier |
|---|---|---|
| Alexa568 anti-rabbit IgG | Life Technologies | A10042; RRID: AB_2534017 |
| IRDye680LT anti-mouse IgG | LI-COR Biosciences, Bad Homburg, Germany | P/N 925-68020; RRID: AB_2687826 |
| IRDye800CW anti-goat IgG | LI-COR Biosciences | P/N 925-32214; RRID: AB_2687553 |
| IRDye800CW anti-mouse IgG | LI-COR Biosciences | P/N 925-32212; RRID: AB_2716622 |
| IRDye800CW anti-rat IgG | LI-COR Biosciences | P/N 925-32219; RRID: AB_2721932 |
| IRDye800CW anti-chicken IgG | LI-COR Biosciences | P/N 925-32218 |
| MFP488 anti-rat IgG | MoBiTec GmbH (Göttingen, Germany) | MFP-A1006 |
| PE-anti-rat IgG (Poly4054) | Biolegend | #405406; RRID: AB_315017 |
| APC Rat IgG1k isotype control | Biolegend | #400411; RRID: AB_326517 |
| FITC Rat IgG1k isotype control | Biolegend | #400405; RRID: AB_326511 |
| PE-rat IgG1k isotype control | Biolegend | #400407; RRID: AB_326513 |

| Substance | Source/Company | Identifier |
|---|---|---|
| FITC-AcLDL | Life Technologies | L23380 |
| Collagen IV | Santa Cruz | sc-29010 |
| Collagenase | Sigma | C2674 |
| Collagenase D | Roche | 11088858001 |
| Dexamethasone | Sigma | D1756 |
| Fluorogold | Life Technologies | H22845 |
| Heparin Natrium 25000 | Ratiopharm | n.d. |
| Histodenz | Sigma | D2158 |
| Insulin | Sigma | I0516 |
| Ketamin | WDT | G2501-04 |
| Mitomycin-C | Pharmacy | n.d. |
| HGF, recombinant | R&D Systems | 294-HG |
| OPN, recombinant | R&D Systems | 441-OP |
| Xylazin | Ecuphor | 400177 |

| Commercial Assays | Source/Company | Identifier |
|---|---|---|
| Mouse Angiogenesis Antibody Array | R&D Systems | ARY015 |
| Quantikine ELISA Hgf | R&D Systems | MHG00 |
| Quantikine ELISA Opn | R&D Systems | MOST00 |
| Gene Chip MOGene ST2.0 | Affymetrix/Thermo Fisher | 902119 |
| GeneChip™ WT Pico Kit | Affymetrix/Thermo Fisher | 902623 |

| Deposited Data | Source | Identifier |
|---|---|---|
| Expression data ECs (wt) | Gene Expression Omnibus (GEO) | GSE 128042 |
| Expression data ECs (wt, early, late) | Gene Expression Omnibus (GEO) | GSE 128044 |
| Expression data SVEC4-10 (ctrl, Hgf 3 h and 6 h) | Gene Expression Omnibus (GEO) | GSE 128046 |

| Cell Lines/Mouse Strains | Source | Species/Origin |
|---|---|---|
| Hep56.1D | Cell lines services GmbH | mouse/hepatocyte |
| SVEC4-10 | ATCC | mouse/endothelial cell |
| HepG2 | ATCC | human/hepatocyte |
| Sk-Hep1 | ATCC | human/endothelial cell |
| LAP-tTA/Yap S127A | kindly provided by Fernando Camargo | |
| AlbTag | kindly provided by Eduard Ryschich | |
| C57Bl/6N | Janvier Labs | |

| Primers for qPCR (m/h) | Genbank ID | Sequence (5'-3') |
|---|---|---|
| Actin (m) | NM_007393 | for: GCTTCTTTGCAGCTCCTTCGT |
| | | rev: ACCAGCGCAGCGATATCG |
| Alb (m) | BC049971.1 | for: GAGGCTGCAAGAAACCTAGGAAGAG |
| | | rev: CACCAGGGATCCACTACAGCAC |
| Bcam (m) | NM_020486 | for: CACGGGAGCACCCTGAGCATTAC |
| | | rev: GAACTCTGAGCCCTGTGGTTCCAC |

| | | |
|---|---|---|
| B2M (h) | NM_004048.3 | for: CACGTCATCCAGCAGAGAAT |
| | | rev: TGCTGCTTACATGTCTCGAT |
| Bmp2 (m) | NM_007553.3 | for: CAACACCGTGCGCAGCTTCC |
| | | rev: GCAGATGTGAGAAACTCGTCAC |
| Cd31 (m) | NM_001032378 | for: GAACCCATCAGGAGTGAATACGTC |
| | | rev: GAACTATGCACCTAATGTGCAGC |
| Cd32b (m) | NM_001077189 | for: GGAGAATATCGGTGTCAAATGGAGC |
| | | rev: CTATGGCACCTTAGCGTGATGG |
| Cd105 (m) | NM_007932 | for: CATGGACAGCCTCTCCTTCCAG |
| | | rev: CCAAGTCCAGATGGCAGCTATCTAG |
| Cd146 (m) | NM_023061.2 | for: GCAACTTCAGCCAAGTGGACTGG |
| | | rev: TCAGTATCTGCCTCTCCTTGTG |
| Clec1b (m) | NM_019985 | for: TCTGCTGATCTCATCCATGGG |
| | | rev: CGCTGAGAGATTTTCCTTCTCCG |
| Ctgf (m) | NM_010217 | for: CAACCGCAAGATCGGAGTGTG |
| | | rev: CCATCCAGGCAAGTGCATTGG |
| CTGF (h) | AY395801.1 | for: CCAAGGACCAAACCGTGG |
| | | rev: CTGCAGGAGGCGTTGTCAT |
| Cxcr4 (m) | NM_009911 | for: GTGATCCTGGTCATGGGTTACCAG |
| | | rev: GAGGTCAGCCACTGACAGGTG |
| Cyr61 (m) | NM_010516 | for: GATCTGTGAAGTGCGTCCTTGTGG |
| | | rev: GACACTGGAGCATCCTGCATAAG |
| Edn1 (m) | NM_010104 | for: CGTGACTTTCCAAGGAGCTCCAG |
| | | rev: CTGAGTTCGGCTCCCAAGAC |
| Epha4 (m) | NM_007936 | for: GCTTGGCTGGAACCAGATCGAC |
| | | rev: GCGATAGCTGCGTTCATTCTGATC |
| Gapdh (m) | NM_008084 | for: TGTCCGTCGTGGATCTGAC |
| | | rev: CCTGCTTCACCACCTTCTTG |
| Hgf (m) | D10213.1 | for: GGTCTGGACTTACATGTTCCATG |
| | | rev: GCTAGCATCTGGCTCCCAGAAG |
| Hprt (m) | NM_013556 | for: TCCTCCTCAGACCGCTTTT |
| | | rev: CCTGGTTCATCATCGCTAATC |
| Lyve-1 (m) | NM_053247 | for: CCACTAGGCACCCAGTCCAA |
| | | rev: GTTTCTGCCCACAAGGGCAAC |
| Met (m) | NM_008591.2 | for: GACATTCTCTTCGGGGTGTT |
| | | rev: GTTGGGTCCGTAAAAATGCT |
| Opn (m) | AF515708.1 | for: CGGATGAATCTGACGAATCTCACC |
| | | rev: CAGAAGCTGGGCAACAGGGATG |
| OPN (h) | J04765.1 | for: GATGGCCGAGGTGATAGTGTGG |
| | | rev: GTGATGTCCTCGTCTGTAGCATCAG |
| Pdgfc (m) | AF286725 | for: GTAGAAGTTGAGGAGCCCAGTGATG |
| | | rev: GTCACAGCATTGTTGAGCAGGTC |
| Plgf (m) | NM_001271705 | for: GCCGATAAAGACAGCCAACATCAC |
| | | rev: CATCTCCACATAGAAATGTGGATCC |
| Stab1 (m) | NM_138672 | for: GAACCCTCCTTCTGCTCTGTGTC |
| | | rev: CTTGGTGTGGATGTCGCAAC |
| Stab2 (m) | NM_138673 | for: GCAAGGAGCAAGCTTCTCCTCG |
| | | rev: GACTGGCACTCCGTCTTGATG |
| TEAD4 (h) | NM_201443.2 | for: TGGAGTTCTCTGCCTTCCTG |

| | | |
|---|---|---|
| | | rev: GGACTGGCCAATGTGCACGA |
| Thbd (m) | NM_009378 | for: GTGTGCTGGCTCCAGCTAG |
| | | rev: GGCTGGCATCGAGGAAGGTC |
| Vcam1 (m) | NM_011693 | for: GAACGAAGTATCCACGTGGACATC |
| | | rev: CCACTGAATTGAATCTCTGGATCC |
| Vegfa (m) | NM_001287056 | for: CCAAAGAAAGACAGAACAAAGCCAG |
| | | rev: CGCTCCAGGATTTAAACCGG |
| Vegfb (m) | BC046303 | for: GTGTGACTGTGCAGCGCTGTG |
| | | rev: GCATTCACATTGGCTGTGTTCTTC |
| Vegfc (m) | NM_009506 | for: CAGCGTAGATGAGCTGATGTCTGTC |
| | | rev: GAAGGTGTTTGTGGCTGCTCC |
| Vegfr3 (m) | BC138348 | for: CACCTGCACCGCCTATGGAG |
| | | rev: GTTCACAGCATCCTGAGTGGTCAC |
| Vwf (m) | NM_011708 | for: CTGCAGTTATCTCCTGGCTGG |
| | | rev: CCCAAGATACACAGACAGGCTC |
| YAP (h) | NM_006106 | for: CCTGCGTAGCCAGTTACCAA |
| | | rev: CCATCTCATCCACACTGTTC |

| **ChIP primers (h)** | **Ensembl ID** | **Sequence (5'-3')** |
|---|---|---|
| OPN downstream control (h) | ENSG00000118785 | for: AGACGGAGTCTCGCTCCGTC |
| | | rev: AATTATCCGGGCGTGGTGGC |
| OPN promoter (h) | ENSG00000118785 | for: ACAAGGATAGGTAGGCTGGGCG |
| | | rev: GTTGCAGAAGTAAAGCAGTTTCTGAC |

| **siRNA** | **Genbank ID** | **Sequence (5'-3')** |
|---|---|---|
| Nonsense | Eurofins Genomics GmbH | UGGUUUACAUGUCGACUAA-dT-dT |
| TEAD4 #1 | NM_201443.2 | GGGCAGACCUCAACACCAA-dT-dT |
| TEAD4 #2 | NM_201443.2 | CCGCCAAAUCUAUGACAAA-dT-dT |

| **Buffer/Solution** | **Ingredients** |
|---|---|
| Amino acid solution (pH 7.6) | 3 mM alanine, 1 mM aspartic acid, 3 mM asparagine, 2mM citrulline, 1 mM cysteine, 6 mM histidine, 6 mM glutamic acid, 13 mM glycine, 3 mM isoleucine, 6 mM leucine, 8 mM lysine, 3.5 mM methionine, 5 mM ornithine, 3 mM phenylalanine, 5 mM proline, 6 mM serine, 1 mM threonine, 3 mM tryptophane, 3 mM tyrosine, 7 mM valine |
| Blocking solution | 5% BSA in TBST |
| Borate buffer (pH 8.8) | 20 mM boric acid, 1.27 mM EDTA |
| $CaCl_2$ Solution | 125 mM |
| Collagen 4 coating solution | 50 mM HCl |
| Collagenase buffer | 64.25% glucose solution, 9.5% HEPES buffer, 9.5% KH buffer, 12,5% amino acid solution, 4% $CaCl_2$ solution, 0.25% glutamine solution |
| Erythrocyte lysis buffer (pH 7.3) | 155 mM $NH_4Cl$, 10 mM $KHCO_3$, 0.1 mM EDTA |
| EGTA solution | 125 mM |
| EGTA buffer | 64.35% glucose solution, 10% KH buffer, 10% HEPES buffer, 15% amino acid solution, 0.4% EGTA solution, 0.25% glutamine solution |
| Glucose solution | 50 mM |
| HEPES buffer | 250 mM HEPES |
| KH buffer | 1 M NaCl, 2 mM KCl, 1 mM $KH_2PO_4$ |
| MACS buffer | 2.5 mM EDTA in PBS, 0.5% BSA |

| | |
|---|---|
| Methocel solution | 1,2% methocel in medium of choice |
| $MgSO_4$ solution | 100 mM |
| Mouse tail lysis buffer (pH 8) | 50 mM tris, 100 mM EDTA, 100 mM NaCl, 1% SDS |
| PBS (pH 7.4) | 140 mM NaCl, 2.7 mM KCl, 10 mM $Na_2HPO_4x2H_2O$, 1.8 mM $KH_2PO_4$ |
| PBST (pH 7.4) | PBS + 0.02% Tween 20 |
| Protein sample buffer (3x, pH 7.6) | 0.01% bromphenol blue, 6% SDS, 15% beta-mercaptoethanole, 30% glycerole, 188 mM tris-HCl |
| Proteinase K solution (pH 8) | 20 µl proteinase K (10 mg/ml), 50 mM tris,10 mM EDTA, 10 mM NaCl |
| RIPA buffer | 150 mM NaCl, 0.1% SDS, 0.5% Na-deoxycholate, 1% Igepal CA 630, 5 mM EDTA, 50 mM tris pH 8, filter sterile |
| SDS running buffer | 25 mM tris, 192 mM glycin, 0.1% SDS |
| Suspension buffer | 63.55% glucose solution, 10% KH buffer, 10% HEPES buffer, 15% amino acid solution, 0.25% glutamine solution, 0.8% $CaCl_2$ solution, 0.4% $MgSO_4$ solution |
| TAE buffer (pH 8) | 40 mM tris acetate, 1 mM EDTA |
| Taliniadis elution buffer | 70 mM tris pH 8, 1 mM EDTA, 1.5% SDS |
| TBST | 0.1% Tween 20 in TBS |
| TE buffer | 70 mM tris pH 8, 1 mM EDTA |
| Tris buffered saline (TBS, pH 7.6) | 20 mM tris-HCl, 140 mM NaCl |

| **Software and algorithms** | **Source** | **Version** |
|---|---|---|
| Aperio | https://www.leicabiosystems.com/digital-pathology/manage/aperio-imagescope/ | Leica Biosystems |
| FlowJo | Analyses performed in the laboratory of Prof. Hanno Glimm / Dr. Claudia Ball | Flowjo FACS analysis |
| FIJI | https://fiji.sc/ | Image Analysis |
| Trackmate | https://imagej.net/TrackMate | FIJI plugin, Trackmate for single cell tracking |
| Linear stack alignment with SIFT | https://imagej.net/Linear_Stack_Alignment_with_SIFT | Plugin for stack registration |
| Skeletonize | https://github.com/fiji/Skeletonize3D | pre-installed function to analyze cellular networks |
| WEKA segmentation | https://imagej.net/Trainable_Weka_Segmentation | pre-installed learning-based segmentation tool |
| Ilastik | https://www.ilastik.org/ | The interactive learning and segmentation toolkit |
| R | https://cran.r-project.org/index.html | R_3.3.3 |
| Ggplot2 | https://ggplot2.tidyverse.org/ | |
| PHATE | https://github.com/KrishnaswamyLab/PHATE | 1.0.0 |
| MAGIC | https://github.com/KrishnaswamyLab/MAGIC | 2.0.3 |
| Complex Heatmap | https://bioconductor.org/packages/release/bioc/html/ComplexHeatmap.html | |
| Circlize | https://cran.r-project.org/web/packages/circlize/index.html | |
| Rcolorbrewer | https://cran.r-project.org/web/packages/RColorBrewer/index.html | |
| EBImage | https://www.bioconductor.org/packages/release/bioc/html/EBImage.html | |

| | | |
|---|---|---|
| RGL | https://cran.r-project.org/web/packages/rgl/index.html | |
| Encode Database | https://www.encodeproject.org/ | Encyclopedia of DNA elements |
| JASPAR | http://jaspar2016.genereg.net/ | JASPAR – a database of transcription factor binding profiles |
| Morpheus | https://software.broadinstitute.org/morpheus/ | Versatile matrix visualization and analysis software |

*Immortalized cell lines and primary liver cells*

The mouse EC line SVEC4-10, the human liver EC line Sk-Hep1, as well as the human liver cancer cell line HepG2 was obtained from the American Type Culture Collection (ATCC; LGC Standards, Wesel, Germany). The murine liver cancer cell line Hep56.1D was obtained from Cell Lines Services GmbH (CLS; Eppelheim, Germany). Cells were cultured in Dulbecco's modified eagle medium (DMEM; SVEC4-10, Sk-Hep1, Hep56.1D) and Roswell Park Memorial Institute medium-1640 (RPMI; HepG2) supplemented with fetal calf serum (FCS, 10%) and 1% penicillin/streptomycin. All cell lines were grown in at 37°C and 5% CO2 in a 95% humidity atmosphere. Cell lines were authenticated by STR-analysis (DSMZ, Braunschweig, Germany) and routinely checked for mycoplasma contamination.

Primary murine ECs and hepatocytes were isolated from 8 - 12 weeks old C57/Bl6N mice. Purified ECs were seeded on collagen IV-coated plastic dishes and cultured using MCDB-131 medium with 10% FCS and 1% penicillin/streptomycin (CCPro, Oberdorla, Germany). Primary hepatocytes were cultivated on collagen I-coated 10 cm dishes (Corning Life Sciences, Amsterdam, Netherlands) in Williams adhesion medium (Biochrom, Berlin, Germany) containing 100 nM dexamethasone, 2 mM glutamin, 10% FCS, and 1% penicillin/streptomycin for 4 hours to support attachment. For the generation of conditioned medium, attached hepatocytes were cultivated in serum free Williams medium supplemented with 2 mM L-glutamin and 1% penicillin streptomycin. Medium was removed after 24 hours for further analysis. Conditioned medium was centrifuged (2,000 x g, 10 minutes), filtered (pore size 0.22 µm, Merck/Millipore, Darmstadt, Germany) and stored at -80°C for further analysis.

*Stimulation Experiments*

Cultured SVEC4-10 cells were starved with serum-free DMEM or RPMI-1640 medium supplemented with 1% penicillin/streptomycin for 3 hours. New serum-free medium containing the growth factors HGF (20 ng/ml) or OPN (200 ng/ml) was administered and cells were further cultured as indicated followed by total mRNA or protein isolation (R&D Systems, Minneapolis, USA).

*siRNA Transfection*

For liposomal transfection of siRNAs with Oligofectamine (Thermo Fisher), cells were seeded into 6 well plates one day prior to transfection. Nonsense siRNA-transfected cells were used as controls. SiRNAs were used at a final concentration of 40 nM (Eurofins Genomics, Ebersberg, Germany) and diluted in Opti-MEM (Gibco/Life Technologies). Cells were harvested for further analysis 72 hours after transfection.

*Mouse models*

C57/Bl6N were bought from Janvier labs (Saint-Berthevin Cedex, France). LAP-tTA/Col1A1-Yap$^{S127A}$ (LAP-Yap) mice were used in this study (1). For transgene repression of constitutively active Yap$^{S127A}$,

mice received 2 mg/ml doxycycline in their drinking water supplemented with 10 mg/ml sucrose. For transgene induction, doxycycline was withdrawn at the age of ten weeks. Control mice (with doxycycline) and animals with $Yap^{S127A}$ expression were sacrificed 8 to 15 weeks after transgene induction.
In general, three to four mice were pooled to obtain sufficient EC numbers for one biological sample. If the liver weight of any individual mouse in one pooled sample exceeded the weight of 5 g, the pooled sample was declared as "late". If all individual liver weighed less than 5 g, the sample was declared as "early". Average liver weights in the "early" and "late" subgroups were 3.29 ± 0.82 g and 5.98 g ± 2.01 g, respectively. Normal liver weight was 1.3 ± 0.2 g.

*Cabozantinib treatment of LAP-tTA/Col1A1-$Yap^{S127A}$ mice*
For cabozantinib treatment, 5 injections (30 mg/kg body weight or carrier solution only (63% NaCl solution (0,9%), 30% polyethylene glycole 300, 5% tween and 2% DMSO) were intraperitoneally injected every other day and livers were explanted 5 days after the last treatment. For the long-term treatment protocol, 4 weekly injections (30 mg/kg body weight) were performed and tissues were collected 1 week after the last treatment.

*Study Approval*
Animal work was authorized by the German Regional Council of Baden-Württemberg (Regierungspräsidium Karlsruhe, ref. numbers G30/13, G201/14, G176/16). All experiments were performed in accordance with the institutional regulations of the IBF (Interfakultäre Biomedizinische Forschungseinrichtung, University of Heidelberg) under pathogen-free (SPF) conditions. The mouse colony was housed under a 12 hours light/dark cycle with free access to water and food. Exclusion and termination criteria were defined in the ATBW criteria.
Human HCCs were surgically resected at the University Hospital and histologically classified according to established criteria by two experienced pathologists. The study was approved by the institutional ethics committee of the Medical Faculty of Heidelberg University (application no. 206/05). Transcriptome and clinical data of 242 human HCC patients have been published previously (2).

*Isolation of Primary ECs and EC Subpopulations*
After median laparotomy, the portal vein was cannulated with a 24 G catheter (Angiocath, BD Biosciences, Heidelberg, Germany) and perfused with collagenase D solution (2 mg/ml; Roche, Grenzach-Whylen, Germany). After liver removal, liver tissue was minced and further digested with collagenase D solution for 45 min at 37°C. Tissue suspension and minced liver tissue were then meshed using cell strainers (100, 70, and 40 µm; BD Biosciences). Prior to gradient centrifugation, remaining erythrocytes in the samples were lysed using erythrocyte lysis buffer (155 mM NH4Cl, 10 mM KHCO3, 0.1 mM EDTA pH 7.3). A non-ionic gradient of Histodenz (Sigma) was used to purify the NPC fraction by a centrifugation step with a disabled break. The NPC fraction was obtained carefully and ECs were magnetically separated using MACS LS columns and LSEC microbeads (Miltenyi Biotec GmbH, Bergisch Gladbach, Germany) according to the manufacturer's instructions. Vital ECs were counted through Trypan blue exclusion in a hemocytometer. Isolated ECs were directly used for cultivation on collagen IV-coated dishes (500,000 cells/cm²), FACS analysis, cell sorting and/or total mRNA extraction.
For the purification of CEC- and LSEC-specific cell fractions, magnetically enriched ECs were FACS stained for CD31, CD146, and Lyve-1 together with respective unstained and isotype controls for FACS gating and compensation. Fluorogold staining (2 µg/ml, Life Technologies, Darmstadt, Germany) was

used to exclude dead cells in the UV excitation channel (excitation wavelength UV laser: 355 nm, detection wavelength 530 ± 30 nm), which allowed the simultaneous exclusion of retinoid-positive HSCs, which emit autofluorescence after UV laser excitation (3). CD31 was detected by APC (excitation wavelength 633 nm, detection wavelength 670 ± 30 nm). FITC (Lyve-1) and PE labeling (CD146) were excited using the blue (488 nm) or yellow/green (561 nm) lasers and detected in the 525 ± 50 nm and in the 575 ± 25 nm range.
Gating of viable double positive CECs (CD31+, CD146+, Lyve-1-, retinoid-), triple-positive LSECs (CD31+, CD146+, Lyve-1+, retinoid-), and cell fraction purification was performed using a cell sorter with a 100 µm nozzle (BD Biosciences, FACSAria, settings for high purity). Total RNA was extracted using the PicoPure extraction kit (Life Technologies) according to the manufacturer's protocol immediately after cell isolation. Flow cytometry was performed using LSRII (analysis) or AriaII (analysis and sorting; Becton Dickinson).

Isolation of Primary Hepatocytes
Primary murine hepatocytes were isolated by continuous liver perfusion with EGTA buffer and collagenase 1 buffer (Sigma) using an ISM596D pump system (Ismatec, Wertheim, Germany). Forceps-based removal of the liver capsule and collection of digested hepatocytes in suspension buffer was followed by filtration through a 100 µm cell strainer (BD Biosciences). Two consecutive low velocity centrifugation steps (50 x g, 2 minutes) were performed to remove the NPC fraction. Pelleted primary hepatocytes were counted and tested for their viability using Trypan blue exclusion. Only samples with vitality >80% were used for cultivation (seeding density: 5x106/10 cm dish). Buffers used for hepatocyte isolation are listed above.

*Spheroid Formation*
SVEC 4-10 spheroids were generated under serum-free conditions using the hanging drop approach (4). The next day, spheroids containing 1,000 cells were embedded in growth factor-reduced Matrigel (Corning Life Sciences) with or without supplementation of HGF (20 ng/ml). Sprouting was digitally documented 48 hours after spheroid embedding.

*Single Cell Migration*
For single cell analysis, 40,000 SVEC4-10 cells were seeded into each 6-well plate. Cells were starved for 3 hours in serum free medium, treated with Mitomycin C to block cell proliferation (2 µg/ml; Pharmacy of the University Hospital Heidelberg) and stained with NucBlue® (Gibco/Life Technologies) according to the manufacturer's instructions. After stimulation with HGF (20 ng/ml), cells were documented for up to 10 hours with 3 measurements/hour using the Olympus Cell Live Cell Imaging System with a IX81 motorized stage, inverted microscope, Hamamatsu camera, and climate chamber (37°C and 5% CO2). Images were acquired using the Olympus Xcellence RT software (Olympus, Hamburg, Germany). The ImageJ plugin "Linear stack alignment with SIFT" was applied to exclude movement artifacts due to incorrect positioning of the motorized stage. Nuclear movement was tracked and measured using the FIJI plugin "Trackmate" (5).

*Data Presentation and Statistics*
Image contrast and brightness were adjusted if needed. In this case, all images within one experiment were adjusted identically. In few cases, a default background subtraction algorithm implemented in ImageJ (rolling ball, default settings) was applied to reduce background signal.

Data is presented as mean ± SD if not stated otherwise. Statistical analysis and graphs were generated using R software, GraphPad Prism, SPSS, and Excel (see table software and algorithms). Labeling of graphs and panel organization were done using Adobe Photoshop and Affinity Designer. In vitro studies were biologically repeated at least three times unless otherwise specified. Statistical analysis was performed as indicated in the figure legends. Mann-Whitney U test was used for the comparison of not normally distributed data. Ordinal data points were associated using the Spearman rank correlation analysis. Unpaired students t-test was used for large patient cohorts with normally distributed data. Multiple groups were compared using Analysis of Variance (ANOVA) followed by post hoc testing. The number of animals used for each experiment are indicated in the figure legends. In vitro and in vivo experimental procedures were compared to respective controls with p values: *$p \leq 0.05$, **$p \leq 0.01$, ***$p \leq 0.001$, ns. not significant.

*Scanning Electron Microscopy (SEM)*

Freshly isolated ECs were cultivated on 18 mm coverslips and fixed in 0.1 M phosphate buffer containing 2% glutaraldehyde. Samples were incubated in 0.1 M cacodylate buffer (pH 7.4) containing 1% OsO4 to increase the contrast of cellular membranes. Dehydration of the samples using acetone was followed by critical point drying. Samples were gold sputtered and microscopy was performed with a Zeiss Leo 1530 SEM. Sample preparation and SEM were performed in collaboration with Ulrike Ganserer and Dr. Ingrid Hausser (Institute of Pathology, Heidelberg and CellNetworks electron microscopy core facility).

*Western Immunoblotting*

For the isolation of total protein fractions from cultured cells, the 10-fold concentrated cell lysis buffer was used (New England Biolabs, Frankfurt, Germany). For the isolation of tissue protein fractions, a small liver piece was homogenized in cell lysis buffer (New England Biolabs) using the Precellys® Ceramic Kit (1.4 mm; 2x 20 seconds, 6,000 rpm; Peqlab, Erlangen, Germany) followed by centrifugation to remove cell debris. Protein concentration was determined using the NanoDrop ND-1000 Spectrophotometer (Thermo Fisher). Equal amounts of proteins were diluted in SDS-PAGE loading buffer (2% SDS, 10% glycerol, 5% β-mercaptoethanole, 0.002% bromphenol blue, 62.5 mM TrisHCl), separated on an 8–12% SDS-PAGE, and electro-transferred to a nitrocellulose membrane in a semidry technique. After blocking of membranes with 5% milk or bovine serum albumin (BSA) in tris-buffered saline/tween 20 (TBST), primary antibodies were added and incubated overnight at 4°C with rotation. The appropriate secondary antibodies were diluted in 5% milk or BSA in TBST (1:20,000; IRDye 680 and 800, LI-COR Biosciences, Bad Homburg, Germany). Membranes were washed 3x with TBST and fluorescence signals were detected using an Odyssey Sa Infrared Imaging System (resolution: 50 µm, offset 3 mm). For normalization, the housekeeping proteins β-actin or GAPDH were detected on each membrane. Antibodies used for Western immunoblotting are listed in Key Resources Table.

*Enzyme-Linked Immunosorbent Assay (ELISA)*

For the detection of soluble factors in cell culture supernatants and mouse blood plasma, ELISAs for murine OPN and HGF were used according to the manufacturer's instructions (R&D Systems). Cytokine concentrations were calculated using respective standard curves. Cell culture supernatants were not diluted, whereas blood plasma samples were diluted 1:10 in dilution buffer.

*Real-time PCR*

Total RNA was isolated using the NucleoSpin® RNA II kit according to the manufacturer's instructions (Macherey Nagel, Macherey-Nagel, Düren, Germany). RNA was eluted in aqua dest. and RNA concentrations were measured at 260 nm using the NanoDrop ND-1000 Spectrophotometer. For cDNA synthesis, 1 µg of total RNA was reversely transcribed using Revert Aid H Minus RT, random hexamer primers and dNTPs (final concentrations 5 µM and 10 mM, respectively) according to the manufacturer's protocol (Thermo Scientific). cDNA was diluted 1:50 for real-time PCR and stored at -20°C.

PCR reactions were set up using the ABsolute qPCR SYBR Green ROX Mix (Thermo Scientific) with the following cycling conditions: 95°C for 15 minutes, followed by 40 cycles of 95°C for 15 seconds, and 60°C for 60 seconds. Melting curve analysis confirmed specific product amplification. For primary cells and tissues, GeNorm was used to normalize the gene expression using the housekeeper genes for mouse Actb, Gapdh, Hprt (https://genorm.cmgg.be). Gene expression of cell lines was normalized using the housekeeper genes for human and mouse β-actin, GAPDH, HPRT, or β2-microglobulin. Samples were measured in technical triplicates. Primers used for semi-quantitative real-time PCR are listed in the Key Resources Table.

*AcLDL Uptake and EC network formation analysis*

The presence of viable ECs in primary liver cell fractions in vitro was confirmed by the EC-specific uptake of acetylated low-density lipoprotein (AcLDL, Thermo Fisher Scientific). For this, ECs were incubated with Al488 conjugated AcLDL according to the manufacturer's recommendations.

For network formation analysis, primary ECs were cultivated for 5 days. IF stains for CD146 and Lyve-1 were performed and fluorescence intensity-based segmentation was performed individually for CD146+ and Lyve-1+ vessel networks. The number of vessel tubes and branches per image were automatically quantified using the ImageJ macro "skeletonize".

*Antibody Array*

For the comparative measurement of pro-angiogenic growth factors and cytokines in the supernatant of starved cultured primary hepatocytes, proteome profiler antibody arrays were used according to the manufacturer's instructions (53 factors on mouse angiogenesis array; R&D Systems). For this, 1 ml of the respective sample was incubated overnight at 4°C on a rocking platform shaker. Bound biotinylated primary antibodies were detected using a DyLight 800 avidin probe and the Odyssey Sa Infrared Imaging System (LI-COR Biosciences). For the comparison of individual membranes, dot blot intensities values were normalized using the positive control dots on each membrane. Hepatocyte derived supernatants of wt (n=3) and transgenic livers (n=5) were analyzed.

*Chromatin immunoprecipitation (ChIP)*

Potential binding sites of TEAD4 in human promoters of the CTGF and SPP1 (gene coding for OPN) were identified using the software tool JASPAR (http://www.jaspar.genereg.net). As negative controls, random sequences downstream of the potential TEAD4 binding sites were chosen and amplified. ChIP was performed as previously described (6). Binding site-specific primers spanned an amplicon of 80-120 bps (listed in Key Resources Table).

*Immunohistochemistry*

Formalin-fixed, paraffin embedded tissue sections were cut into 3 µm thick sections and mounted on microscope slides. Samples were deparaffinized by repeated incubation in xylene (3x for 5 minutes) followed by rehydration in 100% ethanol (2x for 2 minutes), 96% ethanol (1x for 2 minutes) and 70% ethanol (2x for 2 minutes) and rinsed in aqua destillata. Antigen retrieval was achieved by steaming and incubation with target retrieval solution (pH 6, DAKO, Hamburg, Germany). Tissue sections were washed in TBST and blocked with an avidin/biotin blocking kit (Vector Laboratories, Burlingame, CA, USA). The incubation with the primary antibody was performed in a wet chamber at 4°C overnight. After three washing steps with TBST (5 minutes each), either the secondary biotin-conjugated antibody or the Enhancer Detection Line was incubated for 30 minutes (DCS, Hamburg, Germany). Followed by three washing steps in TBST (5 minutes each), the streptavidin horseradish peroxidase (HRP, DAKO) or alkaline phosphatase (AP)-Polymer detection line (DCS) incubation were added, followed by chromogen development (either AP-Red or aminoethylcarbazole (AEC), Zytomed, Berlin, Germany and DAKO, respectively). Antibodies used for immunohistochemistry are listed in Key Resources Table.

*Immunofluorescence and Microscopy*

Snap-frozen tissue pieces were cut to 5 µm thick sections using a Leica CM 3050 S cryostat (Leica Biosystems, Wetzlar, Germany). Tissue sections were fixed in ice-cold acetone for 10 minutes and stored at -20°C. Prior to staining, tissue sections were blocked in PBS containing 10% swine serum at room temperature for 15 minutes. Both, primary and secondary antibodies were incubated at room temperature for 30 minutes. After three final washing steps, slides with tissue sections were dried and mounted with DAPI-containing Fluoromount G. Antibodies used for immunofluorescence are listed in Key Resources Table. Fluorescence microscopy was performed using either an Axiovert 25 microscope (Olympus), a CKX41 (Olympus), or a Confocal DMRE (Leica). For confocal microscopy, laser power and pinhole settings were adjusted prior to image acquisition. All images in one experiment were recorded using identical settings to allow later image quantification. Confocal images were acquired by three-fold detection of the microscopic field to reduce salt and pepper noise.

*Tissue Microarrays*

The HCC tissue microarray used in this study contained 7 non-tumorous liver tissues and 91 HCCs (7x G1, 66x G2, 14x G3, 4x G4 HCCs). For the evaluation of individual immunohistochemical stains both quantity and intensity were evaluated. Quantity was scored as following: 0 = no expression, 1 = up to 1% of cells positive, 2 = 1 to 10% of cells positive, 3 = 11 to 50% of cells positive, 4 = >50% of cells positive. Staining intensity was scored from 0 to 3. 0 = unstained, 1 = weakly, 2 = moderately and 3 = strongly positive. Both, quantitative and qualitative values were multiplied resulting in a score ranging from 0 to 12, which was used for further statistical analysis.

*mRNA Expression Profiling*

Prior to hybridization, RNA integrity was measured (Agilent 2100 Bioanalyzer, Agilent, Frankfurt, Germany). For primary cells, only samples with a RNA integrity number (RIN) >7 were used for transcriptomics. For the generation of CEC and LSEC expression profiles, RNA samples were pre-amplified using the WT Pico Reagent Kit (Affymetrix, High Wycombe, UK). For RNA samples derived from hepatocytes no pre-amplification was needed. Purified and fragmented complementary RNA was generated according to the manufacturer's instructions. Fragments were biotin-labelled prior to

hybridization on MoGene-2_0-st chips using a GeneChip Hybridisation oven 640. Successive staining and scanning were performed with a GeneChipFluidics Station 450 and a GeneChip Scanner 3000, respectively (Thermo Fisher).
After gene annotation, the fluorescence intensity was measured, normalized, and differential expression was statistically assessed using the software package SAS JMP7 genomics (SAS Institute, Cary, NC, USA). As cut-off, a false discovery rate (FDR) value of 0.05 was considered as significant. To assure a homogeneous distribution of the generated data, principal component analysis (PCA) was performed to compare the similarity of individual biological samples in this study. To identify pathways and cellular processes with significant enrichment of differentially expressed genes, gene set enrichment analysis (GSEA) was performed. Enrichment scores were plotted using R studio and the "ggplot2" package. Identified KEGG pathways, including "Cytokine – Cytokine Receptor Interaction – 04060" or "Chemokine signalling – 04062" were used for heatmap generation. Heatmaps were created using the online tool Morpheus (www.software.broadinstitute.org/morpheus) or by R studio and the packages "ComplexHeatmap", "Circlize" and "RColorBrewer". To display expression differences, relative expression differences were displayed by annotating blue and red to the minimum and maximum intensity value for each gene.

*Single Cell Tracking*
Sub-confluent SVEC4-10 cells were stained with NucBlue® (Gibco/Life Technologies) and nuclear movement was tracked using the FIJI plugin "Trackmate" (5). Prior to analysis, image stacks were registered using the ImageJ plugin "Linear stack alignment with SIFT" to exclude movement artifacts through improper positioning of the microscope stage. Individual movement tracks were generated with "Trackmate" and visualized as overlays. In brief, nuclear movement trajectories of intensity-segmented nuclei were generated through an automatized connection of moving nuclei within the image stack using the nearest neighbor algorithm and an upper cut-off value to avoid connection of independent tracks. Tracks were then used to calculate cell-specific individual velocities and distances. For further analysis, nuclear tracks and timepoint-specific velocities were averaged and plotted.

*Measurement of Fluorescence Intensity and Data Analysis*
Fluorescence intensity was displayed by either annotating different colors (16 colors mode) or by generating surface plots (ImageJ/R EBImage and RGL packages). For this, each pixel was depicted in a spatial 3D system with the Z coordinate representing the intensity value. 3D surface plots were either generated by ImageJ using the function "3D surface plots" or by converting the image information into a data matrix, which was then processed in R studio using the RGL package.
For the standardized comparison of Bcam, Thbd, CD31, and CD146 stains, the width of the analyzed field was defined as a line of 10 pixels thickness. The area of interests and their intensity values were exported, averaged, and normalized to LSEC intensities.
For the calculation of EC areas, IF images for CD146 and Lyve-1 were segmented based on their fluorescence intensity using the "Threshold" function in ImageJ.

*Learnable Segmentation Tools*
For an objective analysis of spheroid assays, the segmentation tool "WEKA segmentation" was used (FIJI). In brief, each brightfield image was converted into the three categories "background", "sprouts", and "spheroid core". For each of these image categories, multiple areas of interest were

specified for training of the segmentation algorithm. Once segmentation was sufficient, images were classified. The areas of interest were tagged and measured using the wand tool and measure function.

*ENCODE Database*

The encyclopedia of DNA element (ENCODE) database was used to analyze TEAD4 binding at the SPP1 and CTGF promoters (www.encodeproject.org). TEAD4-dependent sequencing of the SPP1 and CCL2 promoter regions was obtained for the cell lines ECC-1 (GSM accession GSM 1010885) and hESC (GEO accession GSM1010845). ChIP sequencing data were deposited by Richard Myers laboratory, Hudson Alpha Institute for Biotechnology, Huntsville, USA.

*Analysis of scRNA-seq data*

scRNA-seq data from healthy human liver tissues were downloaded from Gene Expression Omnibus (Accession GSE124395), (7). The cell barcodes assigned to clusters 9, 10, 13, 20, 29, and 32 in the original publications were extracted to obtain ECs. Furthermore, the cells were filtered for the expression of PECAM1 (>1). The expression values of obtained ECs were normalized, transformed by applying the square root and visualized using PHATE (8). To emphasize the difference between the groups the expression values used to color the cells were log-transformed.

scRNA-seq data from human HCC patients were downloaded (Set1, 12 patients) from Gene Expression Omnibus (Accession GSE125449), (9). The cell barcodes assigned to cluster TEC in the original publication were extracted to obtain tumor-associated ECs. Furthermore, the cells were filtered to have the imputed expression of PECAM1 greater than 0.75. The tool MAGIC was used to impute missing values (10). The expression values of obtained ECs were normalized, transformed by applying the square root and visualized using PHATE (8). To emphasize difference between groups, the expression values used to color the cells were transformed by taking the $4^{th}$ power. To calculate the correlation between genes for malignant cells, the cell barcodes assigned to cluster "Malignant cell" in the original publication were extracted. The Pearson correlation was calculated using R ($r_P$-values are shown).

**References**


1. Camargo FD, Gokhale S, Johnnidis JB, Fu D, Bell GW, Jaenisch R*, et al.* YAP1 increases organ size and expands undifferentiated progenitor cells. Curr Biol **2007**;17:2054-60
2. Roessler S, Jia HL, Budhu A, Forgues M, Ye QH, Lee JS*, et al.* A unique metastasis gene signature enables prediction of tumor relapse in early-stage hepatocellular carcinoma patients. Cancer Res **2010**;70:10202-12
3. Mederacke I, Dapito DH, Affo S, Uchinami H, Schwabe RF. High-yield and high-purity isolation of hepatic stellate cells from normal and fibrotic mouse livers. Nat Protoc **2015**;10:305-15
4. Muller B, Bovet M, Yin Y, Stichel D, Malz M, Gonzalez-Vallinas M*, et al.* Concomitant expression of far upstream element (FUSE) binding protein (FBP) interacting repressor (FIR) and its splice variants induce migration and invasion of non-small cell lung cancer (NSCLC) cells. J Pathol **2015**;237:390-401
5. Tinevez JY, Perry N, Schindelin J, Hoopes GM, Reynolds GD, Laplantine E*, et al.* TrackMate: An open and extensible platform for single-particle tracking. Methods **2017**;115:80-90
6. Weiler SME, Pinna F, Wolf T, Lutz T, Geldiyev A, Sticht C*, et al.* Induction of Chromosome Instability by Activation of Yes-Associated Protein and Forkhead Box M1 in Liver Cancer. Gastroenterology **2017**;152:2037-51 e22

7. Aizarani N, Saviano A, Sagar, Mailly L, Durand S, Herman JS*, et al.* A human liver cell atlas reveals heterogeneity and epithelial progenitors. Nature **2019**;572:199-204

8. Moon KR, van Dijk D, Wang Z, Gigante S, Burkhardt DB, Chen WS*, et al.* Visualizing structure and transitions in high-dimensional biological data. Nat Biotechnol **2019**;37:1482-92

9. Ma L, Hernandez MO, Zhao Y, Mehta M, Tran B, Kelly M*, et al.* Tumor Cell Biodiversity Drives Microenvironmental Reprogramming in Liver Cancer. Cancer Cell **2019**;36:418-30 e6

10. van Dijk D, Sharma R, Nainys J, Yim K, Kathail P, Carr AJ*, et al.* Recovering Gene Interactions from Single-Cell Data Using Data Diffusion. Cell **2018**;174:716-29 e27
